# Immune elicitation by sensing the conserved signature from phytocytokines and microbes via the *Arabidopsis* MIK2 receptor

**DOI:** 10.1101/2021.01.28.428652

**Authors:** Shuguo Hou, Derui Liu, Shijia Huang, Dexian Luo, Zunyong Liu, Ping Wang, Ruimin Mu, Zhifu Han, Jijie Chai, Libo Shan, Ping He

**Author notes:** These authors contributed equally (Shuguo Hou, Derui Liu, Shijia Huang). **Correspondence:** Shuguo Hou, School of Municipal & Environmental Engineering Shandong Jianzhu University, Ping He, Department of Biochemistry & Biophysics Texas A&M University, Libo Shan, Department of Biochemistry & Biophysics Texas A&M University.

## Abstract

Sessile plants encode a large number of small peptides and cell surface-resident receptor kinases, most of which have unknown functions. Here, we report that the *Arabidopsis* receptor kinase MALE DISCOVERER 1-INTERACTING RECEPTOR-LIKE KINASE 2 (MIK2) recognizes the conserved signature motif of SERINE RICH ENDOGENOUS PEPTIDEs (SCOOPs) from plants as well as proteins present in fungal *Fusarium* spp. and bacterial *Comamonadaceae*, and elicits potent immune responses. SCOOP signature peptides trigger diverse immune and physiological responses in a MIK2-dependent manner with a sub-nanomolar sensitivity and directly bind to the extracellular leucine rich-repeat domain of MIK2 *in vivo* and *in vitro*, indicating that MIK2 is the receptor of SCOOP peptides. Perception of SCOOP peptides induces the association of MIK2 and the coreceptors SOMATIC EMBRYOGENESIS RECEPTOR KINASE 3 (SERK3) and SERK4 and relays the signaling through the cytosolic receptor-like kinases *BOTRYTIS*-INDUCED KINASE 1 (BIK1) and AVRPPHB SUSCEPTIBLE1 (PBS1)-LIKE 1 (PBL1). Our study identified a unique plant receptor that bears a dual recognition capability sensing the conserved peptide motif from phytocytokines and microbial proteins via a convergent signaling relay to ensure a robust immune response.

## INTRODUCTION

Sessile plants have evolved a large number of cell surface-resident receptor proteins to sense and respond to developmental and environmental cues. Many of these receptors are receptor kinases (RKs) with an extracellular domain perceiving the cognate ligands, a transmembrane domain, and a cytoplasmic kinase domain activating downstream signaling ^1–3^. Plant RKs are classified into different subfamilies based on their extracellular domains. Leucine-rich repeat-RKs (LRR-RKs) with a few to more than 30 extracellular LRRs represent the largest subfamily RKs in plants ^2,4^. LRR-RKs are found to perceive plant growth hormone brassinosteroids (BRs), various endogenous and exogenous peptide ligands involved in plant growth, reproduction, cell differentiation, immunity, and beyond ^1,4^, hydrogen peroxide ^5^ and quinone ^6^. Upon the ligand perception, LRR-RKs often heterodimerize with SOMATIC EMBRYOGENESIS RECEPTOR-LIKE KINASE (SERK) subfamily LRR-RKs, e.g., BRASSINOSTEROID INSENSITIVE 1 (BRI1)-ASSOCIATED RECEPTOR KINASE 1 (BAK1)/SERK3 and SERK4, leading to the trans-phosphorylation between the cytoplasmic kinase domains of receptors and SERK coreceptors, and the subsequent activation of the cytoplasmic signaling events ^7,8^.

Some plant LRR-RKs are pattern recognition receptors (PRRs) that recognize pathogen- or microbe-associated molecular patterns (PAMPs or MAMPs) or damage-associated molecular patterns (DAMPs) and initiate pattern-triggered immunity (PTI) ^9–11^. For examples, FLAGELLIN-SENSITIVE 2 (FLS2) and ELONGATION FACTOR-TU (EF-Tu) RECEPTOR (EFR) recognize bacterial flagellin and EF-Tu, respectively ^12,13^; PEP1 RECEPTOR 1 (PEPR1)/PEPR2 and RLK7 recognize plant endogenous peptides PEPTIDE 1 (Pep1), and PAMP-INDUCED SECRETED PEPTIDE 1 (PIP1)/PIP2, respectively ^14–17^. Despite the different origins, MAMPs and DAMPs activate the conserved downstream signaling events, including cytosolic calcium influx, reactive oxygen species (ROS) burst, mitogen-activated protein kinase (MAPK) activation, ethylene production, transcriptional reprogramming, callose deposition, and stomatal closure to prevent pathogen entry ^10,18–20^. It is generally believed that the immune response triggered by DAMPs is to amplify MAMP-activated defense since DAMPs are usually induced upon MAMP perception ^9,18,19^. Some plant secreted peptides, such as SERINE-RICH ENDOGENOUS PEPTIDE 12 (SCOOP12), have been shown to induce typical MAMP/DAMP-triggered immune response, the receptors of which have not been identified ^21^.

The LRR-RK MALE DISCOVERER 1 (MDIS1)-INTERACTING RECEPTOR-LIKE KINASE 2 (MIK2) was initially identified as a component of a receptor heteromer involving in guided pollen tube growth by perceiving the female attractant peptide LURE1 ^22^. However, the physical interaction between MIK2 and LURE1 was not detected ^23^; instead, LURE1 bound directly to another LRR-RK PRK6 ^23^, and the pollen-specific PRK6 was shown to be the receptor of LURE1 family peptides biochemically and genetically ^23–25^. The ligand of MIK2 remains enigmatic. In addition, MIK2 plays a role in response to diverse environmental stresses, including cell wall integrity sensing, salt stress tolerance, and resistance to the soil-borne fungal pathogen *Fusarium oxysporum* ^26–29^. A recent study suggested that MIK2 functions as a PRR perceiving an unknown peptide elicitor from *F. oxysporum* to activate plant immunity ^30^. Here, we provide genetic and biochemical evidence that MIK2 is the bona fide receptor of SCOOP family peptides. SCOOP12 directly binds to the extracellular domain of MIK2 and induces MIK2 dimerization with BAK1/SERK4 coreceptors. Interestingly, SCOOP-LIKE signature motifs are also present in the genomes of a wide range of *Fusarium* spp. and some *Comamonadaceae* bacteria. *Fusarium* SCOOP-LIKE peptides activate the MIK2-BAK1/SERK4-dependent immune responses, pointing to a dual role of MIK2 in the perception of both MAMPs and DAMPs.

## RESULTS

### The kinase domain of MIK2 elicits specific responses

We have previously identified RLK7 as the receptor of the secreted peptides PIP1 and PIP2 ^17^. RLK7 belongs to the subfamily XI of LRR-RKs, including PEPRs and MIK2 (Supplementary Figure 1a). Similar to *RLK7* and *PEPR1*, *MIK2* is transcriptionally upregulated upon treatments of flg22, a 22-amino-acid peptide derived from bacterial flagellin, and elf18, an 18-amino-acid peptide derived from bacterial EF-Tu (Supplementary Figure 1a). To explore the specific function of the extracellular and the cytosolic kinase domains of LRR-RKs, we generated a chimeric receptor consisting of the extracellular domain (ECD) of RLK7 and the transmembrane and cytoplasmic kinase domains (TK) of MIK2 (*RLK7^ECD^-MIK2^TK^*). The chimeric *RLK7^ECD^-MIK2^TK^* and the full-length *RLK7* (*RLK7^ECD^-RLK7^TK^*) under the control of the cauliflower mosaic virus (CaMV) *35S* promoter were transformed into the *rlk7* mutant (Figure 1a).

**Figure 1.**
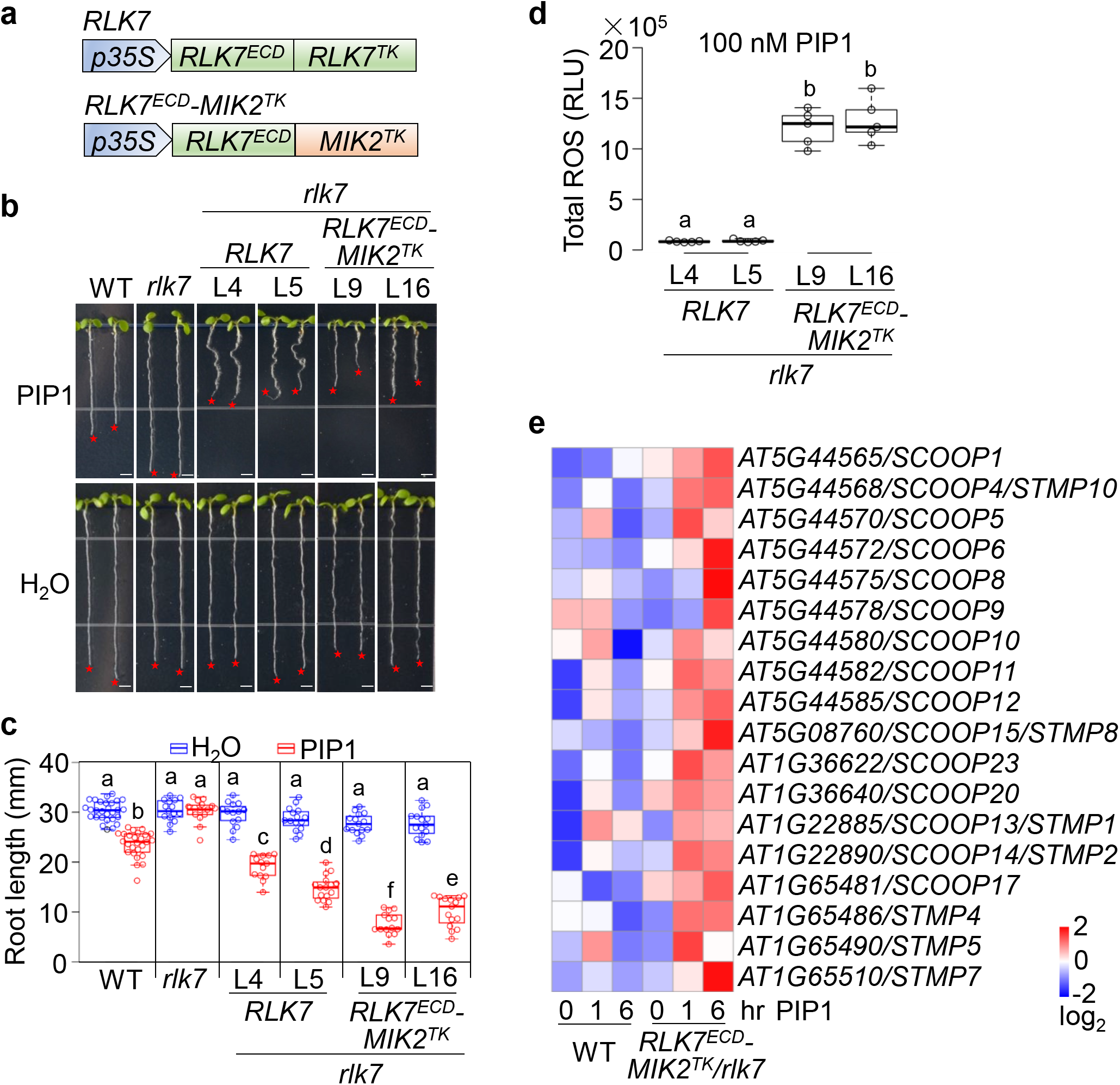
The cytosolic kinase domain of MIK2 induces the expression of *SCOOP* genes. **a.** Diagram of the *RLK7^ECD^-MIK2^TK^* chimeric receptor gene. The extracellular domain of *RLK7* (*RLK7^ECD^*) was fused with the transmembrane and cytoplasmic kinase domain of *MIK2* (*MIK2^TK^*) to generate the *RLK7^ECD^-MIK2^TK^* chimeric gene. The *RLK7^ECD^-MIK2^TK^* chimeric gene under the control of the *35S* promoter was transformed into the *rlk7* mutant. The full-length *RLK7* (*RLK7^ECD^-RLK7^TK^*) driven by the *35S* promoter was also transformed into *rlk7* as a control. **b, c.** *RLK7^ECD^-MIK2^TK^*/*rlk7* transgenic seedlings show more severe root growth inhibition to PIP1 treatment than *RLK7/rlk7* seedlings. Seedlings of *Arabidopsis* WT (Col-0), *rlk7*, two representative lines (L9 and L16) of *RLK7^ECD^-MIK2^TK^*/*rlk7* and two lines (L4 and L5) of *RLK7/rlk7* were grown on ½ MS plates with or without 1 μM PIP1 for ten days (**b**). Red stars indicate the root tips. Scale bar, 4 mm. Quantification data of seedling root length are shown as the overlay of dot plot and means ± SEM (**c**). Different letters indicate a significant difference with others (*P* < 0.05, One-way ANOVA followed by Tukey’s test, *n*=12). **d.** PIP1 treatment induces a more robust ROS production in *RLK7^ECD^-MIK2^TK^*/*rlk7* than that in *RLK7/rlk7* transgenic seedlings. One-week-old seedlings grown on ½ MS plates were treated with 100 nM PIP1, and ROS production was measured immediately with an interval of one second for 15 min. Total luminescence counts were calculated for each treatment. The relative light units (RLU) are shown as the overlay of the dot plot and means ± SEM. Different letters indicate a significant difference with others (*P*< 0.05, One-way ANOVA followed by Tukey’s test, *n*=5). **e.** PIP1 treatment induces the expression of *SCOOP* genes in *RLK7^ECD^*-*MIK2^TK^*/*rlk7* transgenic plants but not in WT plants. Ten-day-old seedlings grown on ½ MS plates were treated with 1 μM PIP1 for 0, 1, and 6 hr for RNA-Seq analysis. Heatmap represents transcript levels of *SCOOPs* and *STMPs.* The original means of gene transcript levels represented by FPKM (Fragments Per Kilobase of exon model per Million mapped fragments) values were subjected to data adjustment by log2 transformation using the OmicStudio (https://www.omicstudio.cn/tool) for the heatmap. The experiments in **b-d** were repeated three times with similar results.

Compared to the *rlk7* mutant, both *RLK7/rlk7* and *RLK7^ECD^-MIK2^TK^/rlk7* transgenic seedlings restored the sensitivity to PIP1 treatment exhibiting root growth retardation (Figures 1b, c), indicating that the chimeric *RLK7^ECD^-MIK2^TK^* receptor is functional. Notably, *RLK7^ECD^-MIK2^TK^/rlk7* seedlings showed more substantial root growth retardation to PIP1 treatment than *RLK7/rlk7* (Figures 1b, c). Besides, PIP1 treatment caused brown roots and darkened root-hypocotyl junctions in *RLK7^ECD^-MIK2^TK^/rlk7* seedlings, but not in *RLK7/rlk7* seedlings (Supplementary Figure 1b). Furthermore, PIP1 treatment triggered a robust production of ROS in *RLK7^ECD^-MIK2^TK^/rlk7* seedlings compared to that in *RLK7/rlk7* seedlings (Figure 1d), which weakly induced ROS production ^17^. We also generated a chimeric receptor carrying the ectodomain of PEPR1 and the transmembrane and cytoplasmic kinase domains of MIK2 (*PEPR1^ECD^-MIK2^TK^*) and transformed it into the *pepr1,2* mutant (Supplementary Figure 1c). The *PEPR1^ECD^-MIK2^TK^/pepr1,2* transgenic seedlings also showed the severe root growth retardation, brown roots, and darkened root-hypocotyl junctions compared to WT seedlings upon Pep1 treatment (Supplementary Figure 1b, d, e). Thus, the MIK2 kinase domain likely triggers specific responses differently from those mediated by the RLK7 and PEPR1 kinase domains.

Interestingly, RNA-sequencing (RNA-seq) analyses revealed that a group of genes encoding secreted peptides of SCOOPs ^21^ and SECRETED TRANSMEMBRANE PEPTIDES (STMPs) ^31^ were explicitly induced at one and/or six hr after PIP1 treatment in *RLK7^ECD^-MIK2^TK^/rlk7*, but not in WT plants (Figure 1e). SCOOPs and STMPs have been recently identified as two groups of secreted peptides with roles in plant growth and defense ^21,31^. SCOOPs and STMPs share certain degrees of similarities at the amino (N)-terminal signal peptide and/or the carboxyl (C)-terminal domain with overlapping family members, such as SCOOP13/STMP1, SCOOP14/STMP2, and SCOOP4/STMP10 (Supplemental Figure 2a).

### Multiple SCOOP peptides activate diverse plant immune and physiological responses

Fourteen *SCOOPs* (*SCOOP1-14*) have been identified in the *Arabidopsis* genome ^21^. Some peptide-encoding genes induced in *RLK7^ECD^-MIK2^TK^/rlk7* seedlings upon PIP1 treatment, such as *AT1G36640*, *AT1G36622*, *STMP4*, *5*, and *7*, bear sequence similarities with *SCOOPs* (Figure 1e). We thus searched the *Arabidopsis* genome and identified additional nine *SCOOP* family members, namely *SCOOP15-23* (Figure 2a, Supplementary Figure 2a). SCOOP15 was also named as STMP8 (Yu et al., 2020). Thus, the *Arabidopsis* genome encodes at least 23 *SCOOP* isoforms characterized by a conserved C-terminus with an SxS motif (where S is serine and x is any amino acid) (Figure 2a, Supplementary Figure 2a) ^21^. Some SCOOPs, including SCOOP6, 7, 10, 11, and 15, contain two copies (an uppercase A or B was added for each copy) of the conserved domain with an SxS motif (Figure 2a). The expression pattern of *SCOOP* genes differs in different plant tissues (Figure 2b). Of those, *SCOOP10* has the highest expression in mature rosette leaves and shoots of seedlings. In contrast, *SCOOP12* and *13* have the highest expression in roots and flowers, respectively (Figure 2b), suggesting that different SCOOPs may bear tissue-specific functions.

**Figure 2.**
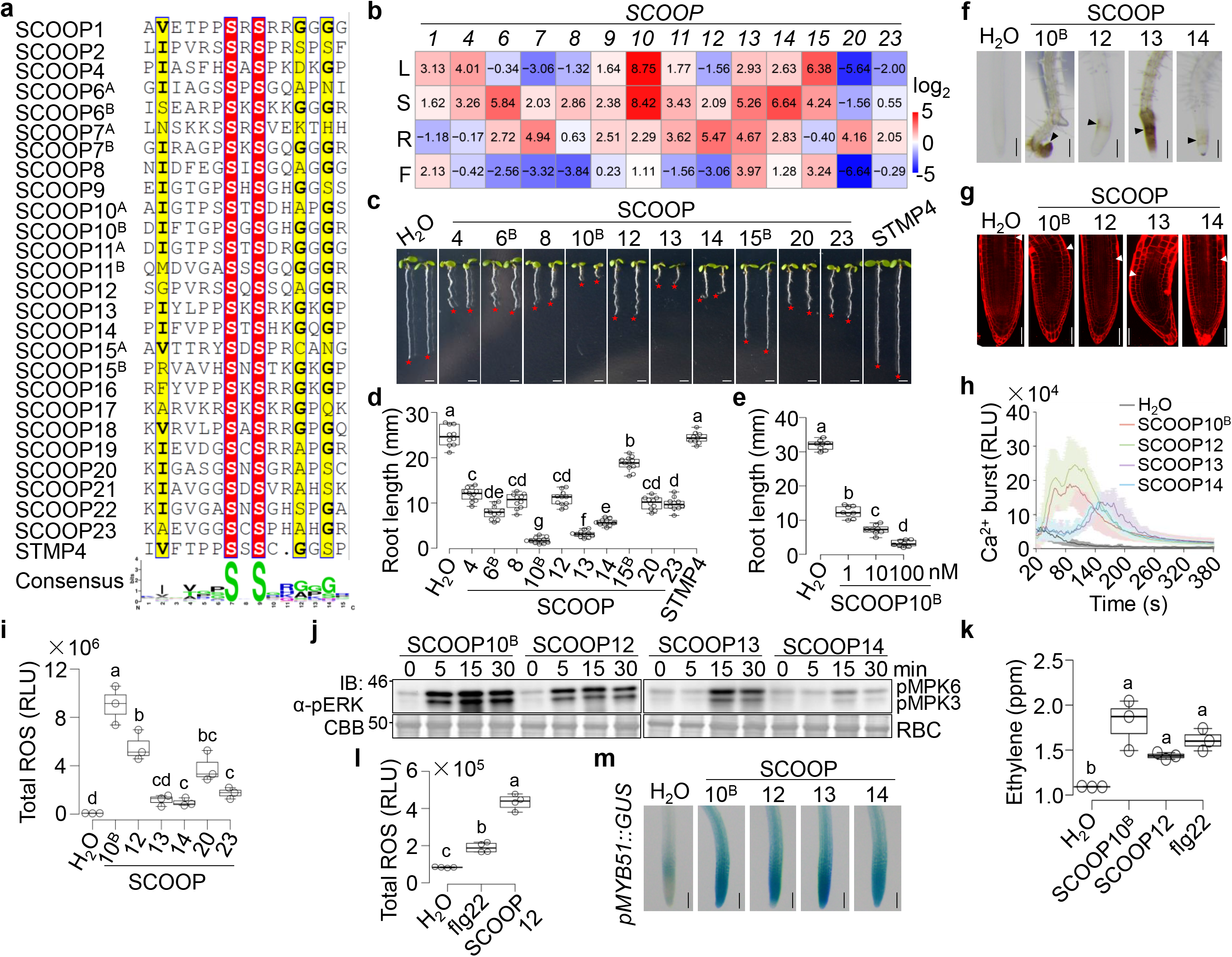
SCOOP peptides activate plant immune responses. **a.** Sequence alignment of SCOOP peptides around the SxS motif. Two domains with the conserved SxS motif in SCOOP6, 10, and 11 are indicated with an uppercase A and B. The amino acid sequences were aligned using ClustalW and displayed in ESPript3 (espript.ibcp.fr). The conserved and similar residues are in red and yellow, respectively. The consensus of SCOOPs was analyzed by WebLogo (http://weblogo.berkeley.edu/logo.cgi). **b.** Expression pattern of *SCOOPs* in different plant tissues. Total RNAs were isolated from the leaves (L), shoots (S), roots (R), and flowers (F) for RT-qPCR analysis of *SCOOPs*. The data represent the average of relative expression levels from three independent repeats with the log_2_ values. **c, d.** SCOOP peptides inhibit root growth. WT seedlings were grown on ½ MS plates with or without 1 μM different peptides for ten days. Quantification data of seedling root length are shown as the overlay of dot plot and means ± SEM (**d**). Different letters indicate a significant difference with others (*P*< 0.05, One-way ANOVA followed by Tukey’s test, *n*≥10). Scale bar, 4 mm. **e.** SCOOP10^B^ inhibits root growth with sub-nanomolar sensitivity. The assay and quantification were performed as in **c, d** with different concentrations of SCOOP10^B^ (*n* ≥ 8). **f.** Treatment of SCOOP peptides causes brown root tips. Root tips of seedlings from (**c**) were photographed under a stereomicroscope. The brown root tips were indicated with an arrowhead. Scale bar, 500 μm. **g.** Treatment of SCOOP peptides distorts root meristems. Root tips of five-day-old WT seedlings grown on ½ MS plates were treated with or without 1 μM peptides for 48 hr followed by propidium iodine (PI) staining and microscopic imaging with a confocal laser scanning microscope. Arrowheads indicate the boundary between the root meristem and the elongation zone. Scale bar, 100 μm. **h.** Treatment of SCOOP peptides triggers cytosolic Ca^2+^ increases. One-week-old transgenic seedlings carrying *p35S::Aequorin* grown on ½ MS plates were treated with or without 100 nM SCOOP peptides, and cytosolic Ca^2+^ concentrations were measured immediately with an interval of one second for 10 min. **i.** Treatment of SCOOP peptides induces ROS production. One-week-old seedlings grown on ½ MS plates were treated with 100 nM peptides. Total luminescence counts as RLU are shown as the overlay of dot plot and means ± SEM. Different letters indicate a significant difference with others (*P*< 0.05, One-way ANOVA followed by Tukey’s test, *n*=5). **j.** SCOOP peptides induce MAPK activation. Ten-day-old WT seedlings grown in ½ MS liquid medium were treated with or without 1 μM peptides for the indicated time. The MAPK activation was analyzed by immunoblotting (IB) with α-pERK antibodies (top panel), and the protein loading is shown by Coomassie brilliant blue (CBB) staining for Rubisco (RBC) (bottom panel). Molecular weight (kDa) was labeled on the left of all immunoblots. **k.** SCOOP peptides induce ethylene production. Leaf discs of four-week-old soil-grown WT plants were incubated in ddH_2_O overnight and then treated with 1 μM peptides for four hr before the measurement of ethylene concentration by gas chromatography. Parts per million (ppm) are shown as the overlay of the dot plot and means ± SEM. Different letters indicate a significant difference with others (*P*< 0.05, One-way ANOVA followed by Tukey’s test, *n*=3). **l.** SCOOP12 peptides induce a more robust ROS production than flg22 in roots. Detached roots from one-week-old WT seedlings grown on ½ MS plates were inoculated in ddH_2_O overnight followed by treatment with 100 nM peptides or ddH2O, ROS production was measured, and data are shown as in i. **m.** SCOOP peptides induce the expression of *pMYB51::GUS* in roots. One-week-old transgenic seedlings carrying *pMYB51::GUS* grown on ½ MS plates were treated with or without 1 μM SCOOP peptides for three hr and subjected to GUS staining followed by photographing under a stereomicroscope. Scale bar, 1 mm. The experiments were repeated three times with similar results.

SCOOP12 has been shown to regulate defense response and root elongation in *Arabidopsis* ^21^. We tested whether other SCOOPs also have the activity to induce immune-related responses in *Arabidopsis*. Based on the sequence characteristics and expression patterns, we synthesized ten SCOOP peptides (SCOOP4, 6^B^, 8, 10^B^, 12, 13, 14, 15^B^, 20, and 23) corresponding to the SxS motif. We also synthesized the STMP4 peptide, which has an amino acid deletion in the conserved SxS motif. Treatment of all synthesized SCOOP peptides, but not STMP4, inhibited root growth to different degrees with SCOOP10^B^ and 13 being the most active (Figures 2c, d). SCOOP10^B^ was highly functional, even on a subnanomole scale (Figure 2e). A close-up view indicated that treatment of SCOOP10^B^, 12, 13, and 14 peptides in WT seedlings led to brown roots, especially at the root tips of seedlings (Figure 2f), and distorted root meristems (Figure 2g), which were not reported for the treatment of flg22 or Pep1 peptides ^14,32^.

Similar to flg22, SCOOP peptides induced various hallmarks of PTI responses, including the cytosolic calcium increase (Figure 2h), ROS production (Figure 2i), MAPK activation (Figure 2j), and ethylene production (Figure 2k). SCOOP10^B^ and 12 exhibited more potent activities than others in triggering these PTI responses (Figures 2h-j). Besides, SCOOP10^B^, 12, 13, and 14 triggered the MAPK activation in both shoots and roots (Supplementary Figure 2b). Notably, SCOOP12-induced ROS production in roots was significantly higher than that elicited by flg22 (Figure 2l). Consistently, SCOOP10^B^, 12, 13, and 14 treatments induced the promoter activities of *MYB51*, a marker gene for root immunity ^33^, in the roots of *pMYB51::GUS* transgenic plants (Figure 2m). The data suggest that SCOOPs play a role in immune activation in both shoots and roots.

### MIK2 mediates SCOOP-induced responses

We further tested if MIK2 is required for SCOOP-triggered responses. We isolated two alleles of *T-DNA* insertional mutants, *mik2-1* (*SALK_061769*) and *mik2-2* (*SALK_046987*). Both *mik2-1* and *mik2-2* were insensitive to the treatment of SCOOP10^B^ (Figures 3a, b). The root growth retardation became more pronounced with the increased concentrations of SCOOP10^B^ from 1 to 100 nM, which was completely blocked in the *mik2-1* mutant (Figures 3c, d). In addition, the *mik2-1* and *mik2-2* mutants were insensitive to the growth inhibition triggered by the other nine SCOOPs (Figure 3e and Supplementary Figure 3a). However, the *mik2-1* mutant responded to the Pep1 treatment similarly to WT plants (Supplementary Figure 3b). Besides, *MIK2-LIKE* (*MIK2L*), the closest homolog of *MIK2*, and other members of subfamily XI LRR-RKs, including *RLK7* ^17^, *PERP1* ^15^, *PERP2* ^16^, *HAESA* (*HAE*), and *HAESA-LIKE2* (*HSL2*) ^34^, were not involved in responsiveness to SCOOP10^B^ (Supplementary Figure 3c). The *fls2* mutant responded to SCOOP10^B^ similarly to WT plants (Supplementary Figure 3c), supporting that the activities in synthesized SCOOP peptides were unlikely due to the contamination of flg22. The data indicate that the root growth inhibition triggered by SCOOP peptides genetically depends on MIK2, but not other members of the subfamily XI LRR-RKs.

**Figure 3.**
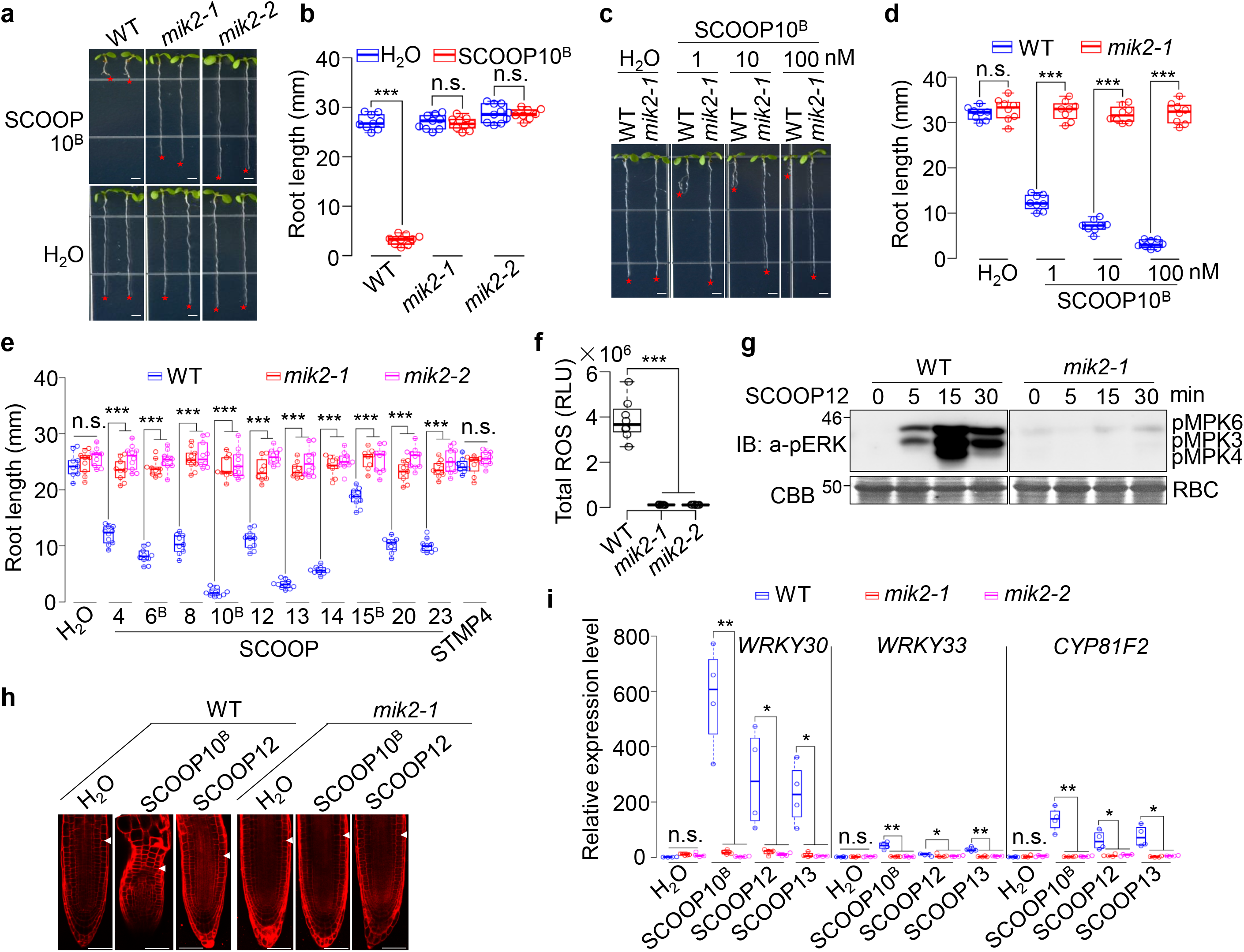
SCOOP-triggered responses depend on MIK2. **a-d.** SCOOP10^B^-triggered root growth inhibition is blocked in *mik2* mutants. WT and *mik2* mutant seedlings were grown on ½ MS plates with or without 1 μM SCOOP10^B^ (**a**) or different concentrations of SCOOP10^B^ (**c**) for ten days. Quantification data of seedling root length are shown as the overlay of dot plot and means ± SEM (**b, d**) (*** *P*<0.001, n.s., no significant differences, Student’s *t*-test, *n*≥8). Scale bar, 4 mm. **e.** The *mik2* mutants block the root growth inhibition induced by different SCOOPs. The assay and quantification were performed as in **a**and **b** with 1 μM of SCOOP peptides. **f.** SCOOP12-induced ROS production is blocked in *mik2* mutants. One-week-old seedlings grown on ½ MS plates were treated with 100 nM SCOOP12. Total luminescence counts as RLU are shown as the overlay of dot plot and means ± SEM (*** *P*<0.001, Student’s *t*-test, *n*=4). **g.** SCOOP12-induced MAPK activation is blocked in the *mik2-1* mutant. Ten-day-old seedlings grown in ½ MS liquid medium were treated with or without 1 μM SCOOP12 for the indicated time. The MAPK activation was analyzed by immunoblotting with α-pERK antibodies (top panel), and the protein loading is shown by CBB staining for RBC (bottom panel). **h.** The *mik2-1* mutant blocks SCOOP-induced root meristem distortion. Root tips of five-day-old seedlings grown on ½ MS plates were treated with or without 1 μM peptides for 48 hr followed by PI staining and microscopic imaging with a confocal laser scanning microscope. Arrowheads indicate the boundary between the root meristem and the elongation zone. Scale bar, 50 μm. **i.** SCOOP-induced defense gene expression is blocked in *mik2* mutants. Ten-day-old seedlings grown on ½ MS plates were treated with or without 1 μM SCOOP10^B^, 12, or 13 for 1 hr and subjected to RT-qPCR analysis. The expression of genes was normalized to that of *UBQ10,* and the relative expression levels are shown as the overlay of the dot plot and means ± SEM (\**P*<0.05, ** *P*<0.01, n.s., no significant differences, Student’s *t*-test, *n*=4). The experiments in **a-d**, **f-i**were repeated three times, and **e** twice with similar results.

We next determined whether SCOOP-mediated other physiological and immune responses also depended on MIK2. The SCOOP12-induced ROS production (Figure 3f) and MAPK activation (Figure 3g) were abolished in the *mik2* mutants. The SCOOP10^B^- and 12-induced distortion of meristems in root tips was not observed in the *mik2-1* mutant (Figure 3h). Reverse transcription-quantitative polymerase chain reactions (RT-qPCR) showed that the induction of immune-related marker genes, *WRKY30*, *WRKY33*, and *CYP81F2*, by SCOOP10^B^, 12, and 13 in WT seedlings was blocked in the *mik2* mutants (Figure 3i). Taken together, these data indicate that SCOOP-triggered responses depend on MIK2.

### MIK2 is the receptor of SCOOP12

Since SCOOPs trigger MIK2-dependent responses, we investigated whether SCOOP12 could bind to MIK2. We synthesized the red fluorescent tetramethylrhodamine (TAMRA)-labeled SCOOP12 peptides at its N-terminus (TAMRA-SCOOP12) and determined the ability of WT and *mik2* plants for binding to TAMRA-SCOOP12 *in vivo*. TAMRA-SCOOP12 peptides were bioactive as they triggered a MIK2-dependent root growth inhibition and MAPK activation, similar to SCOOP12 (Supplementary Figures 4a-c). Red fluorescent signals were detected in roots and leaf protoplasts in WT plants, but not in the *mik2-1* mutant upon treatment of TAMRA-SCOOP12 (Figure 4a). Importantly, pretreatment of unlabeled SCOOP12, but not flg22, markedly reduced red fluorescent signals of TAMRA-SCOOP12 in WT seedlings (Figure 4b), indicating specific and MIK2-dependent binding of SCOOP12 *in vivo*.

**Figure 4.**
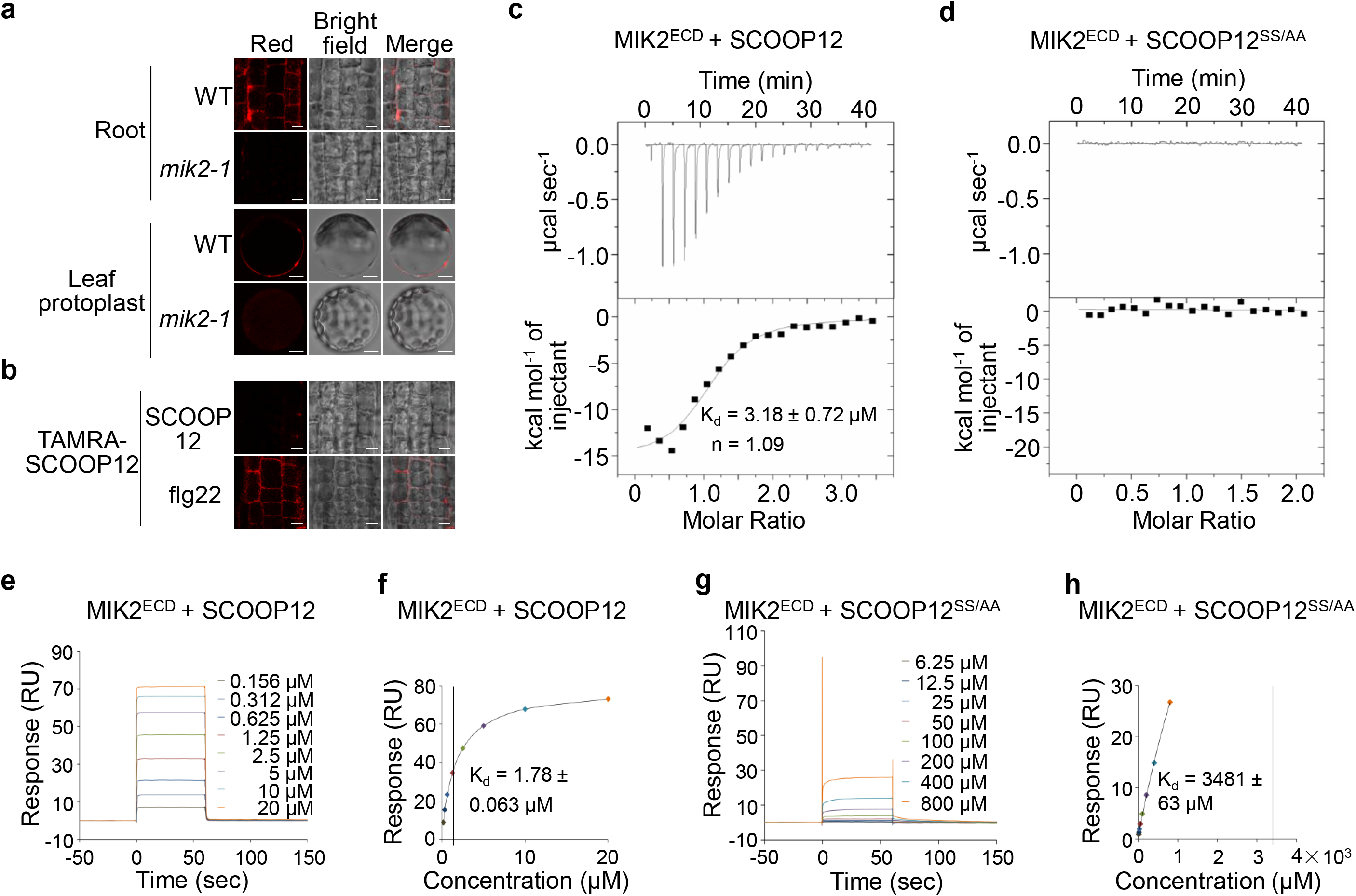
SCOOP12 binds to MIK2. **a.** TAMRA-SCOOP12 peptides label roots and protoplasts from WT but not the *mik2-1* mutant. Five-day-old WT seedlings grown on ½ MS plates were incubated with 100 nM TAMRA-SCOOP12 for 5 min and washed with ½ MS liquid medium three times. Protoplasts from four-week-old leaves were incubated with 100 nM TAMRA-SCOOP12 for 5 min and washed with W5 solution three times. Samples were observed under a confocal laser scanning microscope with 552 nm excitation: scale bar, 4 mm (top), 10 μm (bottom). **b.** SCOOP12 but not flg22 peptides compete for the binding of TAMRA-SCOOP12 to roots. Five-day-old WT seedlings grown on ½ MS plates were incubated with 100 nM TAMRA-SCOOP12 in the presence of 1 μM SCOOP12 or flg22 peptides for 5 min, washed and imaged as in a. **c, d.** SCOOP12 but not SCOOP12^SS/AA^ binds to MIK2^ECD^ with isothermal titration calorimetry (ITC) assays. SCOOP12 (**c**) or SCOOP12^SS/AA^ (**d**) was titrated into a solution containing MIK2^ECD^ in ITC cells. The top panels show raw data curves, and the bottom panels show the fitted integrated ITC data curve. The calculated binding kinetic constant (K_d_ values ± fitting errors) for SCOOP12 with MIK2^ECD^ is 3.18 ± 0.72 μM, and the stoichiometry of binding (n) is approximately equal to one ligand molecule per receptor molecule (**c**). No binding was detected for SCOOP12^SS/AA^ with MIK2^ECD^ **(d)**. **e, f.** SCOOP12 binds to MIK2^ECD^ with surface plasmon resonance (SPR) assays. Affinity-purified MIK2^ECD^ proteins isolated from insect cells were immobilized by an amine-coupling reaction on a sensor chip and synthesized SCOOP12 peptides were used as flow-through analyte for SPR assays. (**e**) shows the SPR sensorgram profile of SCOOP12 peptides at gradient concentrations flowing through the MIK2^ECD^ immobilized chip. (**f**) shows the steady-state affinity (binding at equilibrium) indicated by a calculated K_d_ of 1.78 μM. **g, h.** SCOOP12^SS^/^AA^ does not bind to MIK2^ECD^ with SPR assays. Similar assays using SCOOP12^SS^/^AA^ peptides were performed as in **e**and **f**. The SPR sensorgram profile (**g**) indicated nearly no binding between MIK2^ECD^ and SCOOP12^SS^/^AA^ with a K_d_ of 3481 μM (**h**). The above experiments were repeated twice with similar results.

To test whether SCOOP12 directly binds to MIK2 *in vitro*, we employed isothermal titration calorimetry (ITC) analysis using the extracellular LRR domain of MIK2 (MIK2^ECD^) expressed from insect cells. ITC quantitatively measures the binding equilibrium by determining the thermodynamic properties of protein-protein interaction in solution. As shown in Figure 4c, MIK2^ECD^ bound SCOOP12 potently with a dissociation constant (K_d_) of 3.18 μM. Alanine (A) substitutions on two conserved serine (S) residues in the SxS motif of SCOOP12 (SCOOP12^SS^/^AA^) abolished its activities to inhibit root growth through MIK2 (Supplementary Figure 4d, e) (Gully et al., 2019). No binding was detected between MIK2^ECD^ and SCOOP12^SS^/^AA^ (Figure 4d). The data indicate that MIK2 recognizes and binds to SCOOP12 directly, and the conserved SxS motif is essential for SCOOP12 binding to MIK2. In addition, we performed surface plasmon resonance (SPR) assays, in which affinity-purified MIK2^ECD^ proteins isolated from insect cells were immobilized by an amine-coupling reaction on a sensor chip, and synthesized SCOOP12 peptides were used as the flow-through analyte. The SPR sensorgram showed the profile of SCOOP12 peptides at gradient concentrations flowing through MIK2^ECD^ peptides immobilized on the chip (Figure 4e). The analysis of the binding at equilibrium for SCOOP12 and MIK2^ECD^ indicated a calculated K_d_ of 1.78 μM (Figure 4f), suggesting a high affinity of MIK2^ECD^ with SCOOP12. In contrast, a similar analysis of the SPR sensorgram using the MIK2^ECD^ sensorchip flowing through with SCOOP12^SS^/^AA^ peptides (Figure 4g) indicated nearly no binding between MIK2^ECD^ and SCOOP12^SS^/^AA^ with a K_d_ of 3481 μM (Figure 4h). Together, our data suggest that the extracellular domain of MIK2 directly binds to SCOOP12 with a considerably high affinity, and the conserved SxS motif of SCOOP12 is critical for its binding to MIK2.

### BAK1 and SERK4 are coreceptors for MIK2 in mediating SCOOP-triggered immunity

Ligand perception by LRR-RK receptors often recruits the BAK1/SERK4 family coreceptors to activate downstream signaling ^7,8^. Indeed, SCOOP10^B^- and 12-induced root growth inhibition was partially compromised in *bak1-4* compared to WT (Figures 5a, b). Consistent with the redundant functions of BAK1 and SERK4, the SCOOP-mediated root growth inhibition was alleviated in the *bak1-5*/*serk4* mutant (Figures 5a, b). Similarly, SCOOP12-induced ROS production was reduced in *bak1-4* and completely abolished in *bak1-5*/*serk4* (Figure 5c). In addition, SCOOP12- or 13-induced MAPK activation was reduced in *bak1-4* and substantially blocked in *bak1-5*/*serk4* (Figure 5d, Supplementary Figure 5a). To examine whether the SCOOP perception induces the MIK2 and BAK1 association, we performed co-immunoprecipitation (co-IP) assays in the *mik2-1* mutant transformed with *GFP*-tagged *MIK2* under the control of its native promoter (*pMIK2::MIK2-GFP/mik2*) with α-BAK1 antibodies. The association between MIK2 and BAK1 was barely detectable in the absence of peptide treatment but was stimulated upon the treatment of SCOOP10^B^ or 12 (Figure 5e). SCOOP10^B^- and 12-induced MIK2-BAK1 association was also observed in protoplasts co-expressing the hemagglutinin (HA)-tagged MIK2 (MIK2-HA) and FLAG epitope-tagged BAK1 (BAK1-FLAG) (Supplementary Figure 5b). SERK4 is also associated with MIK2 after SCOOP peptide treatments (Figure 5f).

**Figure 5.**
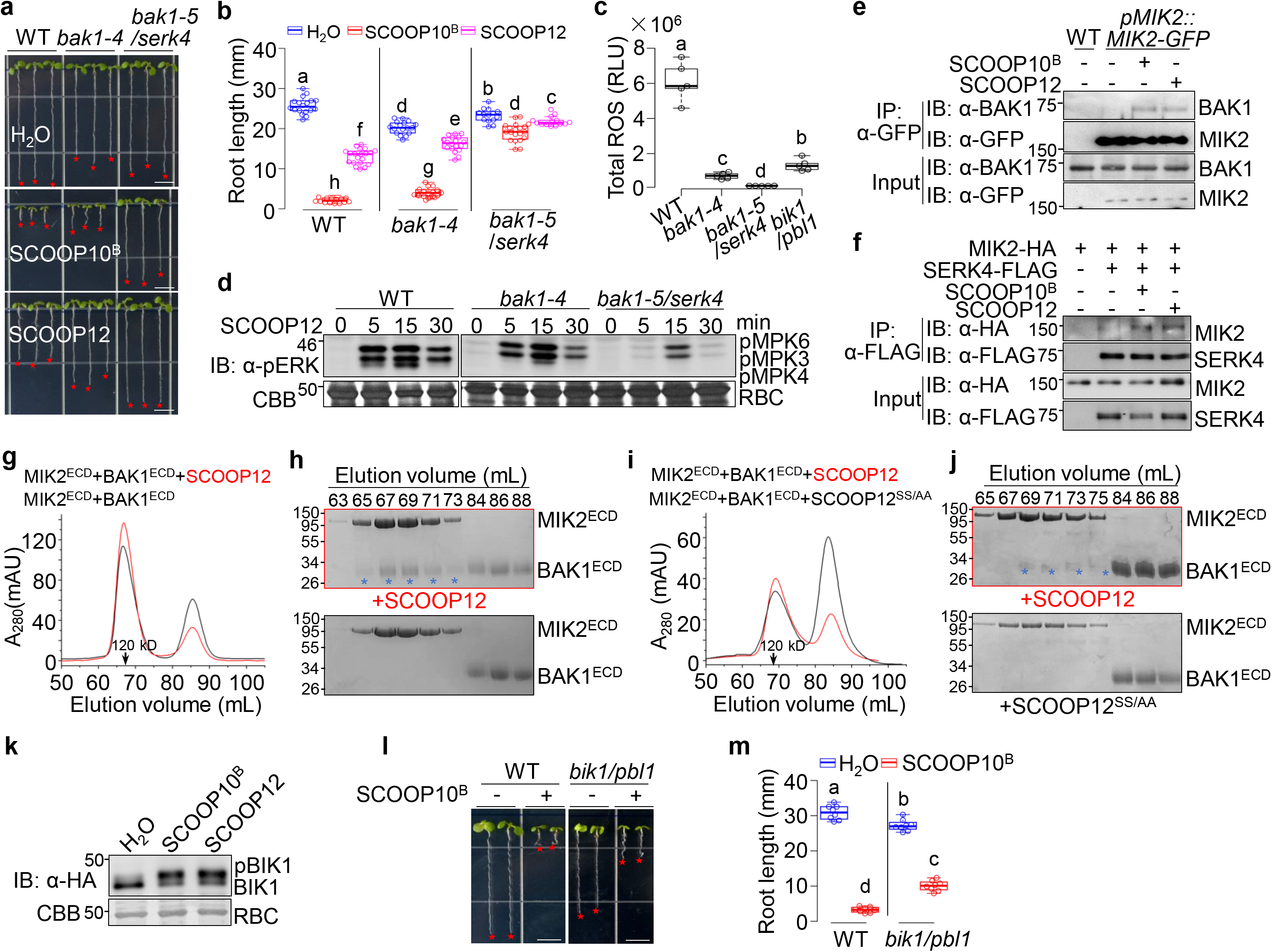
SCOOPs induce the MIK2 and BAK1/SERK4 complex formation. **a, b.** SCOOP10^B^- and SCOOP12-triggered root growth inhibition is compromised in *bak1-4* and *bak1-5/serk4* mutants. Seedlings were grown on ½ MS plates with or without 1 μM SCOOP10^B^ or SCOOP12 for ten days (**a**). Quantification data of seedling root length are shown as the overlay of dot plot and means ± SEM (**b**). Different letters indicate a significant difference with others (*P*<0.05, One-way ANOVA followed by Tukey’s test, *n*≥15). Scale bar, 4 mm. **c.** SCOOP12-induced ROS production is compromised in *bak1-4*, *bak1-5/serk4*, and *bik1*/*pbl1.* One-week-old seedlings grown on ½ MS plates were treated with 100 nM SCOOP12 for ROS measurement with a duration of 15 min. Total luminescence counts as RLUs are shown as the overlay of dot plot and means ± SEM. Different letters indicate a significant difference with others (*P*<0.05, One-way ANOVA followed by Tukey’s test, *n*=5). **d.** SCOOP12-induced MAPK activation is compromised in *bak1-4* and *bak1-5/serk4*. Ten-day-old seedlings grown in ½ MS liquid medium were treated with or without 1 μM SCOOP12 for the indicated time. The MAPK activation was analyzed by immunoblots with α-pERK antibodies (top panel), and the protein loading is shown by CBB staining for RBC (bottom panel). **e.** SCOOPs induce the association of MIK2 and BAK1 in transgenic plants. Leaves of four-week-old transgenic plants carrying *pMIK2::MIK2-GFP* in the *mik2-1* background were treated with or without 1 μM SCOOP^B^ or SCOOP12 peptides for 30 min. Total proteins were subjected for immunoprecipitation (IP) with α-GFP agarose beads (IP: α-GFP), and the immunoprecipitated proteins were detected with α-BAK1 or α-GFP antibodies (top two panels). The input controls before immunoprecipitation are shown on the bottom two panels. **f.** SCOOPs induce the association of MIK2 and SERK4 in protoplasts. Protoplasts from leaves of four-week-old WT plants were co-transfected with HA-tagged MIK2 (MIK2-HA) and FLAG-tagged SERK4 (SERK4-FLAG), or a control vector (Ctrl) and incubated for 10 hr followed by treatment with or without 1 μM SCOOP^B^ or SCOOP12 peptides for 15 min. The IP was performed as in **e**. **g, h.** SCOOP12 induces the interaction between MIK2^ECD^ and BAK1^ECD^. Gel filtration chromatography analysis using MIK2^ECD^ and BAK1^ECD^ isolated from insect cells shows the elution profiles in the presence (red) or absence (black) of SCOOP12 (**g**). SDS-PAGE and CBB staining show the eluted fractions (top panel with SCOOP12; bottom panel without SCOOP12). Stars indicate the eluted BAK1^ECD^ complexing with MIK2^ECD^ (**h**). **i, j.** SCOOP12^SS/AA^ does not induce the interaction between MIK2^ECD^ and BAK1^ECD^. Similar assays were done as above in the presence of SCOOP12 (red) or SCOOP12^SS/AA^ (black). **k.** SCOOP10^B^ and SCOOP12 induce BIK1 phosphorylation. Protoplasts from WT plants were transfected with HA-tagged BIK1 (BIK1-HA) and incubated for 8 hr followed by treatment with 1 μM SCOOP10^B^ or SCOOP12 for 15 min. Proteins were subjected to IB using α-HA antibodies (top panel), and CBB staining for RBC as loading controls (bottom panel). Phosphorylated BIK1 (pBIK1) was indicated as a band mobility shift in IB. **l, m.** SCOOP10^B^-triggered root growth inhibition is compromised in *bik1/pbl1*. Seedlings were grown on ½ MS plates with or without 1 μM SCOOP10^B^ for ten days (**l**). Quantification data of seedling root length are shown as the overlay of dot plot and means ± SEM (**m**). Different letters indicate a significant difference with others (*P*<0.05, One-way ANOVA followed by Tukey’s test, *n*≥8). Scale bar, 4 mm. The experiments in **a-d, l,** and **m** were repeated three times and **e-j** twice with similar results.

To test whether the extracellular domains of MIK2 and BAK1 are sufficient to form a SCOOP12-induced complex *in vitro*, we performed a gel filtration assay of BAK1^ECD^ and MIK2^ECD^ purified from insect cells in the presence of SCOOP12 or SCOOP12^SS/AA^ peptides. The results show that MIK2^ECD^ and BAK1^ECD^ proteins co-migrated in the presence of SCOOP12 (Figures 5g, h), indicating that SCOOP12 induces the dimerization between MIK2^ECD^ and BAK1^ECD^. The protein complex was eluted mainly at the position corresponding to a size of a monomeric MIK2^ECD^-BAK1^ECD^ (~120 KD) (Figures 5g, h), suggesting that SCOOP12 may not induce the homodimerization of the MIK2^ECD^-BAK1^ECD^ complex. In contrast, SCOOP12^SS/AA^, which is unable to bind to MIK2^ECD^, cannot induce the dimerization between MIK2^ECD^ and BAK1^ECD^ (Figures 5i, j), indicating that the SCOOP12-MIK2 binding is required for the MIK2-BAK1 interaction.

### SCOOP-MIK2-BAK1 relays the signaling through the BIK1 family RLCKs

MAMP/DAMP perception activates receptor-like cytoplasmic kinases (RLCKs), including *BOTRYTIS*-INDUCED KINASE 1 (BIK1) and its close homolog AVRPPHB SUSCEPTIBLE1 (PBS1)-LIKE 1 (PBL1) ^35,36^. SCOOP10^B^ or 12 treatments led to the phosphorylation of BIK1, as indicated by a protein band mobility shift in the immunoblot (Figure 5k). SCOOP12-induced ROS burst and SCOOP10^B^-induced growth inhibition were significantly compromised in the *bik1*/*pbl1* mutant (Figures 5c, l, m). Thus, the MIK2-BAK1 receptor complex activation by SCOOPs triggers phosphorylation of BIK1 and PBL1, which further transduces signaling to downstream events. The activation of NADPH oxidase RBOHD by BIK1 is required for PRR-induced ROS production ^37,38^. Likewise, SCOOP12-induced ROS production was abolished in the *rbohD* mutant (Supplementary Figure 5c). Our results collectively indicate that the SCOOP-MIK2-BAK1 receptorsome participates the conserved signaling pathways shared by other MAMP/DAMP/PRR receptor complexes.

### Microbial SCOOP-LIKE peptides trigger a MIK2-dependent immune response

A recent report showed that the *mik2* mutants were insensitive to an unidentified proteinous elicitor isolated from several *Fusarium* spp., suggesting that MIK2 may perceive a peptide elicitor from *Fusarium* ^30^. This prompted us to examine whether SCOOP-LIKE (SCOOPL) peptides are encoded in the genomes of *Fusarium* spp. We blast-searched the genome of *F. graminearum*, a *Fusarium* strain with a well-annotated genome sequence ^39^, using SCOOP10^B^ and 12 as queries. A 21-amino-acid peptide sequence in the N-terminus of an uncharacterized protein (FGSG_07177) showed a high similarity to SCOOP12 (Figure 6a, Supplementary Figures 6a-c). FGSG_07177 is predicted as a transcription factor with a GAL4-like DNA-binding domain (Supplementary Figure 6b). Structural modeling indicates that the SCOOPL region is located on the surface of FGSG_07177 (Figure 6b). Further blast-searching with other available *Fusarium* genome sequences revealed that the SCOOPL domains are highly conserved among all *Fusarium* species surveyed (Figure 6a, Supplementary Figure 6c). Alignment of 22 SCOOPL sequences from different *Fusarium* species revealed seven groups, most of which, except for *F. pseudograminearum*, contain an SxS motif as *Arabidopsis* SCOOPs (Figure 6a, Supplemental Figure 6c). The second conserved serine residue of *F. pseudograminearum* SCOOPL (*Fpg*SCOOPL) is substituted by a proline (Figure 6a, Supplemental Figure 6c).

**Figure 6.**
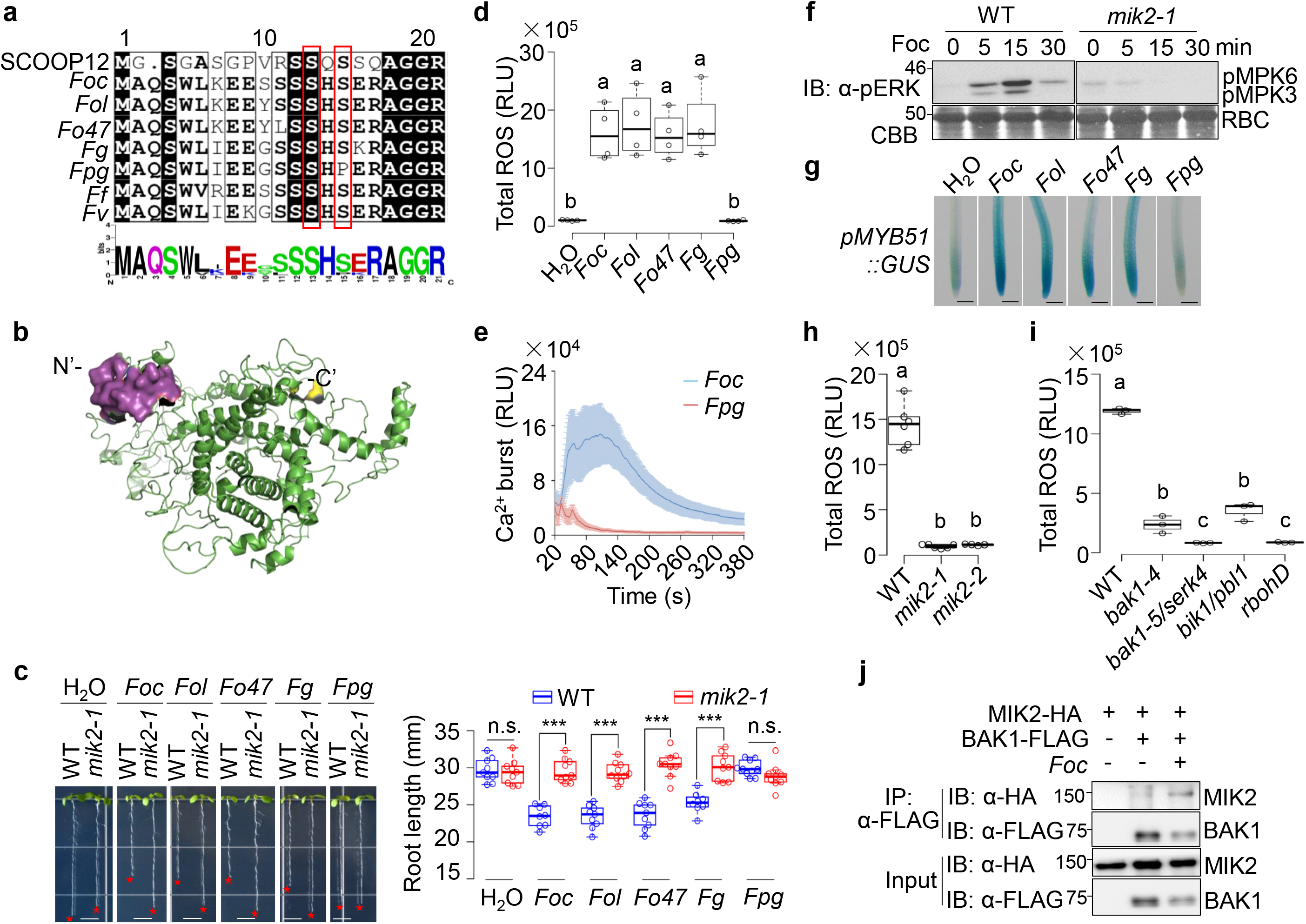
*Fusarium* SCOOP-like peptides activate MIK2-dependent immune responses. **a.** Alignment of SCOOP-LIKE (SCOOPL) sequences around the SxS motif from different *Fusarium* strains using *Arabidopsis* SCOOP12 as the reference. The amino acid sequences were aligned using ClustalW and visualized in ESPript3 (espript.ibcp.fr). The consensus of SCOOPs was conducted with WebLogo (http://weblogo.berkeley.edu/logo.cgi). *Foc*, *F. oxysporum* f. sp. *conglutinans* race 2; *Fol*, *F. oxysporum* f. sp. *lycopersici* MN25, *Fo*47, *F. oxysporum* 47; *Fg*, *F. graminearum* PH-1; *Fpg*, *F. pseudograminearum*; *Ff*, *F. fujikuroi*; *Fv*, *F. venenatum*. Two conserved serine (S) residues are boxed in red. **b.** The surface accessibility of the *Fg*SCOOPL peptide. The FGSG_07177 protein was homology-modeled using Pymol with the centromere DNA-binding protein complex CBF3 subunit B as the template (PDB-ID, c6f07B). The N-terminal 21-amino acid *Fg*SCOOPL peptide is highlighted in magenta, the C-terminus in yellow, and the rest in green. **c.** Multiple *Fusarium* SCOOPL peptides, except *Fpg*SCOOPL, inhibit root growth in a MIK2-dependent manner. WT and *mik2-1* seedlings were grown on ½ MS plates with or without 1 μM *Fusarium* SCOOPL peptides for ten days (left). Quantification data of seedling root length are shown as the overlay of dot plot and means ± SEM (right) (*** *P*<0.001, n.s., no significant differences, two-sided two-tailed Student’s *t*-test, *n*≥8). Scale bar, 4 mm. **d.** Treatment of *Fusarium* SCOOPLs, except *Fpg* SCOOPL, induces ROS production. One-week-old WT seedlings grown on ½ MS plates were treated with or without 1 μM peptides for ROS measurement with a duration of 15 min. Total luminescence counts as RLUs are shown as the overlay of dot plot and means ± SEM. Different letters indicate a significant difference with others (*P*<0.05, One-way ANOVA followed by Tukey’s test, *n*=4). **e.** Treatment of *Foc* SCOOPL triggers the cytosolic Ca^2+^ increase. One-week-old transgenic seedlings carrying *p35S::Aequorin* grown on ½ MS plates were treated with or without 1 μM *Foc*SCOOPL or *Fpg*SCOOPL, and the cytosolic calcium concentrations presented as RLUs were measured immediately for 6 min. **f.** *Foc*SCOOPL-induced MAPK activation is blocked in *mik2-1*. Ten-day-old seedlings grown in ½ MS liquid medium were treated with or without 1 μM *Foc*SCOOPL for the indicated time. The MAPK activation was analyzed by immunoblotting with α-pERK antibodies (top panel), and the protein loading is shown by CBB staining for RBC (bottom panel). **g.** *Fusarium* SCOOPLs induce the expression of *pMYB51::GUS* in roots. One-week-old transgenic seedlings carrying *pMYB51::GUS* grown on ½ MS plates were treated with or without 1 μM SCOOPL peptides for 3 hr and subjected to GUS staining followed by photographing under a stereomicroscope. **h, i.** *Foc*SCOOPL-induced ROS production is blocked in *mik2, bak1-4*, *bak1-5/serk4*, *bik1*/*pbl1*, and *rbohD* mutants. The assay and quantification were performed as in **d**. **j.** *Foc*SCOOPL induces the association of MIK2 and BAK1 in protoplasts. Protoplasts from WT plants were co-transfected with HA-tagged MIK2 (MIK2-HA) and FLAG-tagged BAK1 (BAK1-FLAG), or a control vector (Ctrl), and incubated for 10 hr, followed by treatment with or without 1 μM *Foc*SCOOPL for 15 min. Proteins were immunoprecipitated with α-FLAG agarose beads (IP: α-FLAG), followed by immunoblotting with α-HA or α-FLAG antibodies (top two panels). The input control before IP is shown by IB on the bottom two panels. The experiments in **c-j** were repeated three times with similar results.

To examine whether *Fusarium* SCOOPL peptides trigger immune responses as *Arabidopsis* SCOOPs, we synthesized five *Fusarium* SCOOPL peptides from *F. oxysporum* f. sp. *conglutinans* (*Foc*), *F. oxysporum* f. sp. *Lycopersici* (*Fol*), *F. oxysporum* strain Fo47 (*Fo*47), *F. graminearum* (*Fg*), and *F. pseudograminearum* (*Fpg*). Four out of five *Fusarium* SCOOPLs, except *Fpg*SCOOPL, which has a polymorphism in the second conserved serine, were able to induce growth inhibition (Figure 6c), ROS production (Figures 6d), Ca^2+^ influx (Figure 6e, Supplementary Figures 7a, b), MAPK activation (Figure 6f), and *MYB51* promoter activation in roots (Figure 6g). *Fg*SCOOPL and *Foc*SCOOPL induced the cytosolic Ca^2+^ increase at the concentration of 10 nM and 100 nM, respectively (Supplementary Figures 7a, b). *Fusarium* SCOOPL appeared to trigger a weaker ROS production than *Arabidopsis* SCOOP10^B^ and SCOOP12 (Supplementary Figure 7c). Notably, the *Fusarium* SCOOPL-induced growth inhibition, MAPK activation, and ROS production were abolished in the *mik2* mutant (Figures 6c, f, h). Moreover, similar to SCOOP12, the *Foc*SCOOPL-induced ROS production was compromised in the *bak1-5*/*serk4*, *bik1*/*pbl1*, and *rbohD* mutants (Figure 6i). *Foc*SCOOPL also induced the MIK2-BAK1 association in *Arabidopsis* protoplasts (Figure 6j). Thus, *Fusarium* SCOOPLs trigger similar responses with *Arabidopsis* SCOOPs in a MIK2-dependent manner. This is consistent with the observation that the *mik2* mutants were insensitive to the elicitor from different *Fusarium* spp. ^30^. Interestingly, SCOOPLs also exist in the C-terminus of an unknown protein conserved in *Comamonadaceae* bacteria, including *Acidovorax*, *Curvibacter* sp., and *Verminephrobacter eiseniae* (Supplemental Figure 7d). SCOOPLs from *Curvibacter* sp. (*Cu*) and *Verminephrobacter eiseniae* (*Ve*), but not from *Acidovorax temperans* (*At*) or *Acidovorax avenae* (*Aa*) induced growth inhibition and *MYB51* promoter activation in roots (Supplementary Figures 7e-g). The first conserved serine residue is absent in *Aa*SCOOPL (Supplementary Figures 7d). The *Cu*SCOOPL- or *Ve*SCOOPL-induced growth inhibition was blocked in the *mik2-1* mutant (Supplementary Figure 7e, f). Thus, SCOOPL signature motifs are highly conserved in different microbes, and they may serve as MAMPs perceived by the plant MIK2 receptor.

## DISCUSSION

The *Arabidopsis* genome encodes more than 1000 putative small peptide genes, most of which have unknown functions ^40^. Similarly, plants have also evolved a large number of RKs, with only a few having defined ligands and functions ^3^. In this study, we report that LRR-RK MIK2 specifically recognizes multiple plant endogenous peptides of SCOOP family members, leading to a series of PTI responses, including cytosolic Ca^2+^ influx, ROS burst, MAPK activation, ethylene production, and defense-related gene expression. SCOOP12 directly binds to the extracellular LRR domain of MIK2 *in vivo* and *in vitro*, indicating that MIK2 is a *bona fide* receptor of SCOOPs. The SxS signature motif is essential for SCOOP functions and MIK2 binding. Perception of SCOOPs by MIK2 induces the heterodimerization of MIK2 with BAK1 and SERK4, and BAK1/SERK4 are required for SCOOP-triggered responses, indicating that BAK1/SERK4 are co-receptors of MIK2 in perceiving SCOOPs. Interestingly, the SCOOP active motif was also detected in a conserved, but uncharacterized protein ubiquitously present in fungal pathogen *Fusarium* spp. and bacterial *Comamonadaceae*. Both *Fusarium* and *Comamonadaceae* SCOOP-LIKE peptides with the SxS signature activate the MIK2-BAK1/SERK4-dependent immune responses. Thus, our data reveal a dual role of MIK2 in perceiving the conserved SCOOP signature motif from plants and pathogens in plant immune activation.

Plant plasma membrane-resident RKs perceive diverse exogenous and endogenous signals via the extracellular domain and activate intracellular responses through the cytoplasmic kinase domain ^1–3^. The chimeric receptors with the swapped extracellular and intracellular domains between different RKs have been used to study the specificity of signal perception and signaling activation ^41–46^. The extracellular domains of RKs determine the ligand-binding specificity ^43,44,47^. The intracellular kinase domains of different RKs also trigger specific responses ^41,45^. For example, the chimeric receptor of the EFR extracellular domain and wall-associated kinase 1 (WAK1) cytoplasmic kinase domain triggers defense responses that are activated by oligogalacturonides (OGs), the proposed ligand of WAK1 ^41^. We show that the chimeric receptor of RLK7^ECD^-MIK2^TK^ or PEPR1^ECD^-MIK2^TK^ activates some immune responses that are usually not observed upon RLK7 or PERP1 activation by the corresponding ligands. More importantly, the immune responses triggered by RLK7^ECD^-MIK2^TK^ or PEPR1^ECD^-MIK2^TK^ mirror those usually activated by SCOOPs, the ligands of MIK2. For example, RLK7 activation by ligand PIP1 moderately inhibits root growth and weakly induces ROS production ^17^. In contrast, the RLK7^ECD^-MIK2^TK^ activation by PIP1 causes severe growth inhibition and robust ROS production, similar to SCOOP treatments. Coincidentally, PIP1-activated responses are BIK1-indenependent ^17^, whereas SCOOPs induce BIK1 phosphorylation and BIK1-dependent responses. Thus, upon ligand perception by the extracellular domain, the cytosolic kinase domain of RKs activates some convergent and unique signaling events, likely through differential phosphorylation events, and interaction with different partners.

Fourteen *SCOOPs* have been previously identified in *Arabidopsis*, and SCOOP12 can activate defense response and regulate root elongation ^21^. Our study extends the SCOOP family to 23 members, and all tested ones with the conserved SxS motif activate plant immune response. It remains unknown why plants have evolved so many SCOOPs and the functional specificity for different SCOOPs. Notably, *SCOOP* genes show different expression patterns with some highly expressed leaves and some in roots. *MIK2* is highly expressed in leaves and roots (http://bar.utoronto.ca/efp/cgi-bin/efpWeb.cgi, and data not shown). Thus, it is likely that specific SCOOPs might be recognized by MIK2 in different plant tissues. *SCOOP12*, one of the most active SCOOPs, is highly expressed in roots. Coincidentally, SCOOP12 triggers a robust immune response in roots, and physiological changes, such as darkened hypocotyl-root junctions and distorted meristems in roots. In addition, the *mik2* mutant shows increased susceptibility to the root-invading pathogen *F. oxysporum* ^26^. However, the enhanced susceptibility to *F. oxysporum* has not been observed in the *scoop12* mutant (data not shown), likely due to the functional redundancy of different *SCOOPs*.

Surprisingly, SCOOPL sequences are also present in a wide range of the fungal *Fusarium* spp. and bacterial *Comamonadaceae*. In the fungal *Fusarium* spp, the SCOOPL motif is located in the N-terminus of a highly conserved GAL4 DNA-binding domain-containing protein with uncharacterized functions. It remains unknown whether this protein is secreted during *Fusarium* infection and how MIK2 perceives it. Notably, elf18, an 18-amino acid peptide perceived by LRR-RK EFR, is derived from the conserved bacterial translation elongation factor EF-Tu, which is unlikely to be a secreted protein ^48^. It will also be interesting to determine whether *Fusarium* SCOOPL is the proteinous elicitor isolated from several *Fusarium* spp., which triggers a MIK2-dependent immunity ^30^. Some SCOOPL peptides from *Fusarium* spp. and *Comamonadaceae* have amino acid variations in the conserved SxS motif and cannot activate the immune response. This is consistent with the observation that the SxS motif is required for plant SCOOP peptide functions and binding to MIK2. The variations may be due to the pathogens’ evolutionary pressure to escape from hosts’ perception of immune elicitation.

It has been reported that some pathogens and nematodes deploy mimics of plant endogenous peptides, such as CLAVATA3/ESR (CLE), PLANT PEPTIDE CONTAINING SULFATED TYROSINE (PSY) and RAPID ALKALINIZATION FACTOR (RALF), to promote the pathogenicity by hijacking plant peptide-receptor signaling ^49–52^. For example, RALF secreted from *F. oxysporum* induces plant receptor FERONIA-dependent extracellular alkalization to favor fungal multiplication ^49^. In contrast, similar to plant SCOOPs, microbial SCOOPLs activate the MIK2-dependent immune responses, suggesting that SCOOPLs act as MAMPs rather than virulence factors. Compared to the wide distribution of SCOOPLs in fungal *Fusarium* spp. and bacterial *Comamonadaceae*, plant SCOOPs are only present in *Brassicaceae* genus ^21,31^. This indicates that plant *SCOOPs* may have evolved later than microbial *SCOOPLs*. In addition, gene duplications of *SCOOPs* are common in *Brassicaceae* species ^21^, suggesting that *SCOOPs* are highly evolved genes. For example, twelve *SCOOPs* exist as tandem repeats on *Arabidopsis* chromosome 5 ^21^. Thus, plant SCOOPs may have evolved to mimic microbial SCOOPL and amplify SCOOPL-triggered immunity. Moreover, plant SCOOPs are generated from peptide precursor proteins, whereas *Fusarium* SCOOPLs reside in the N-terminus of a putative transcription factor. The divergence of two precursor protein classes suggests that SCOOPs and SCOOPLs might have evolved convergently but unlikely by horizontal gene transfers ^53^.

## MATERIALS AND METHODS

### Plant materials and growth conditions

The *Arabidopsis thaliana* accession Columbia-0 (Col-0) was used as wild-type (WT). T-DNA insertion mutants of *mik2-1* (*SALK_061769*), *mik2-2* (*SALK_046987*), *mik2-like* (*SALK_112341*) were obtained from the Nottingham *Arabidopsis* Stock Centre (NASC). The *fls2*, *bak1-4*, *bik1/pbl1*, *bak1-5/serk4*, *rlk7*, and *rbohd* mutants were described previously ^17,35,54^. The *hae*/*hsl2* mutant was kindly provided by Dr. Reidunn B. Aalen (University of Oslo, Norway), the *pepr1-2/pepr2-2* (*pepr1,2*) mutant by Dr. Zhi Qi (Inner Mongolia University, China), a transgenic line expressing *p35S::Aequorin* by Dr. Marc Knight (Durham University, UK), and transgenic line expressing *pMYB51::GUS* by Dr. Frederick M. Ausubel (Harvard Medical School, US). Plants were grown in soil (Metro Mix 366, Sunshine LP5 or Sunshine LC1, Jolly Gardener C/20 or C/GP) in a growth room at 20-23°C, 50% humidity, and 75-100 μE m^−2^ s^−1^ light with a 12-hr light/12-hr dark photoperiod for 4-5 weeks before protoplast isolation, or ethylene measurement. Seedlings used for analyses of root growth inhibition, MAPK activation, ROS production, cytosolic Ca^2+^ concentration increase, gene transcription, and GUS staining were grown on half-strength Murashige and Skoog (½ MS) plates containing 0.5% (w/v) sucrose, 0.75% (w/v) agar, and 2.5 mM MES, pH 5.8, under the same conditions as plants grown in soil.

### Plasmid construction and generation of transgenic plants

The *pHBT-35S::BAK1-FLAG*, *pHBT*-*35S::SERK4-FLAG*, *pHBT*-*35S::BIK1-HA*, and *pUC19-35S::MIK2-HA* constructs were described previously ^22,35,54^. The fusion protein vector carrying *MIK2^ECD^* or *BAK1^ECD^* for insect cell expression was reported previously ^23,54^. To obtain the binary vector *pCAMBIA1300-35S::RLK7*, a 2907-bp *RLK7* coding sequence (CDS) was PCR-amplified from WT cDNA using gene-specific primers with *Kpn*I and *Sal*I at the 5’ and 3’ ends respectively, followed by *Kpn*I and *Sal*I digestion and ligation into the *pCAMBIA1300* vector. To generate the binary vector *pCAMBIA1300* carrying the *RLK7^ECD^*-*MIK2^TK^* chimeric receptor gene, an 1824-bp fragment encoding *RLK7^ECD^* was PCR-amplified from WT cDNA using a gene-specific forward primer with *Kpn*I at the 5’ end and a gene-specific reverse primer with 8-bp overlapping sequence from the 5’ end of *MIK2^TK^*, and a 1011-bp fragment encoding *MIK2^TK^* using a gene-specific forward primer with 8-bp overlapping sequence from the 3’ end of *RLK7 ^ECD^* and a gene-specific reverse primer with *Sal*I at the 5’ end. *RLK7^ECD^* was fused with *MIK2^TK^* and inserted into the *pUC19* vector using in-fusion recombinant enzymes (Clontech). After digestion with *Kpn*I and *Sal*I, the *RLK7^ECD^*-*MIK2^TK^* chimeric receptor gene was inserted into a *pCAMBIA1300* vector with the *CaMV 35S* promoter and the HA tag at the 3’ end to generate *pCAMBIA1300-p35S::RLK7^ECD^*-*MIK2^TK^*. A similar strategy was used to generate *pCAMBIA1300-35S::PEPR1^ECD^*-*MIK2^TK^*. A 2307-bp fragment encoding *PEPR1^ECD^* was amplified for fusion with *MIK2^TK^*. To obtain *pHBT*-*35S::MIK2-GFP* constructs, the *MIK2* CDS was PCR-amplified from WT cDNA using gene-specific primers with *Bam*HI and *Stu*I at 5’ and 3’ ends, respectively, followed by *Bam*HI and *Stu*I digestion and ligation into the *pHBT* vector with *HA* or *GFP* sequence at the 3’ end. To generate the binary vector *pCAMBIA1300-pMIK2::MIK2-GFP*, a 2000-bp promoter sequence upstream of the start codon of *MIK2* was PCR-amplified and subcloned into *pHBT*-*35S::MIK2-HA* between *Xho*I and *Bam*HI sites. The *pMIK2::MIK2-GFP-NOS* fragment was further PCR amplified and inserted into *pCAMBIA1300* between *Eco*RI and *Sal*I using in-fusion recombinant enzymes. The primers used for cloning and sequencing were described in Supplementary Table 1, and the Sanger-sequencing verified all insertions in different vectors.

The binary vectors were introduced into *Agrobacterium tumefaciens* strain GV3101 for the floral dipping method-based *Arabidopsis* transformation. Transgenic plants were selected with 20 μg/L hygromycin B. Multiple transgenic lines in the T_1_ generation were analyzed by immunoblotting for protein expression. Two lines with a 3:1 segregation ratio for hygromycin resistance in the T_2_ generation were selected to obtain homozygous seeds.

### Peptide synthesis

PIP1, Pep1, flg22, and all non-labeled SCOOPs and SCOOP-LIKE peptides were synthesized at ChinaPeptides (Shanghai, China). TAMRA-labeled SCOOP12 peptides were synthesized at Biomatik (Delaware, USA). The sequences of synthesized peptides were listed in Supplementary Table 2.

### Root growth assay

Cold stratified seeds were surface-sterilized with 70% (v/v) ethanol for 5 min and were sown on ½ MS plates with or without the peptides at the indicated concentrations. Ten-day-old seedlings grown on plates vertically in a growth chamber were photographed, and the root lengths of seedlings were measured using Image J (http://rsb.info.nih.gov/ij/).

### RNA sequencing (RNA-Seq)

Ten-day-old WT Col-0 and *RLK^ECD^-MIK^TK^* (L16) seedlings grown on ½ MS plates were incubated in 1 mL of liquid ½ MS medium overnight. Seedlings were then treated with or without 1 μM PIP1 for 1 or 6 hr and harvested for RNA isolation. Total RNAs (5 μg) from two biological replicates were pooled for cDNA library construction. cDNA library preparation and sequencing were carried out on an Illumina HiSeq 4000 platform with 150-nucleotide pair-end reads in LC-BIO (Hangzhou, China). The raw sequence data have been submitted to the NCBI database with accession number GSE159580. Total reads were mapped to the Arabidopsis genome (TAIR10; www.arabidopsis.org) with Hisat, and the read counts for every gene were generated StringTie. Differentially expressed genes (DEGs) between different treatments were defined by fold change of read counts ≥ 2 with *P-value* ≤ 0.01. PIP1-regulated DEGs in 1 or 6 hr were listed in supplementary table 3.

### RNA isolation and reverse transcription-quantitative polymerase chain reactions (RT-qPCR) analysis

Total RNA was extracted from ten-day-old seedlings grown on ½ MS plates, or from rosette leaves or inflorescences of four- or seven-week-old soil-grown plants using TRIzol reagent (Invitrogen). One microgram of total RNA was reverse-transcribed to synthesize the first-strand cDNA with M-MuLV Reverse Transcriptases (Thermo Fisher Scientific) and oligo(dT) primers following by RNase-free DNase I (Thermo Fisher Scientific) treatment. RT-qPCR analyses were performed on a QuantStudio™ 3 Real-Time PCR Detection System (Thermo Fisher Scientific) using Faster Universal SYBR^®^ Green Master (Roche) and gene-specific primers following the standard protocol. The expression of each gene was normalized to the expression of *UBQ10*. The primers used for RT-qPCR were listed in Supplementary Table 1.

### ROS assay

ROS burst was determined by a luminol-based assay. Five one-week-old seedlings grown on ½ MS plates were incubated in 200 μL ddH2O overnight in a 1.5 mL centrifuge tube. Then, ddH2O was replaced by 200 μL of reaction solution containing 50 μM of luminol, and 10 μg/mL of horseradish peroxidase (Sigma-Aldrich) supplemented with or without 100 nM or 1 μM peptide. Luminescence was measured immediately after adding the solution with a luminometer (Glomax 20/20n, Promega) with a one-second interval for 15 min. The total values of ROS production were indicated as means of the relative light units (RLU).

### Measurement of cytosolic Ca^2+^ concentration

One-week-old seedlings expressing *p35S::Aequorin* grown vertically on ½ MS plates were put into a 1.5 mL centrifuge tube containing 200 μL solution with 1 mM KCl and 1 mM CaCl_2_. Acquorin was reconstituted by treating the seedlings with coelenterazine h (Promega, Beijing, China) in dark overnight at a final concentration of 10 μM. Luminescence was measured with a luminometer (Glomax 20/20n, Promega) with a one-second interval for 10 min. The values for cytosolic Ca^2+^ concentration were indicated as means of relative light units (RLU).

### Measurement of ethylene production

Twelve leaves of four-week-old plants grown on soil were excised into leaf discs of 0.25 cm^2^, followed by overnight incubation in a 25 mL glass vial with 2 mL of ddH2O for recovery. Then, ddH2O was replaced by 1 mL of peptides at 1 μM, and the vials were capped immediately with a rubber stopper and incubated at 23 °C with gentle agitation for 4 hr. One mL of the vial headspace was injected into a TRACE^TM^ 1310 Gas Chromatograph (Thermo Fisher Scientific) with FID supported by Chromeleon 7 for quantitation.

### Histochemical detection of GUS activity

Two-week-old seedlings grown on ½ MS plates were immersed and vacuumed in the GUS staining solution (10 mM EDTA, 0.01% [v/v] Silwet L-77, 2 mM potassium ferricyanide, 2 mM potassium ferrocyanide and 2 mM 5-Bromo-4-chloro-3-indolyl-β-D-glucuronic acid in 50 mM sodium phosphate buffer, pH 7.0) for 5 min, followed by incubation at 37 °C for 12-24 hr. Stained samples were fixed with a 3:1 ethanol:acetic acid solution overnight and cleared by lactic acid and photographed with the Olympus SZX10 stereoscopic microscope.

### Microscopy assays

For the observation of propidium iodide (PI) stained roots, five-day-old seedlings grown on ½ MS plates were mounted in ddH_2_O containing 10 μM PI for 20 min before imaged with the Leica SP8 confocal microscope. Images were captured at 543 nm laser excitation and 578-700 nm emission.

For the observation of TAMRA-SCOOP12 labeled roots and protoplasts, five-day-old seedlings grown on ½ MS plates or protoplasts isolated from leaves of four-week-old soil-grown plants were treated with 100 nM TAMRA-SCOOP12 with or without 1 μM SCOOP12 or flg22 for 5 min in liquid ½ MS or W5 solution (154 mM NaCl, 125 mM CaCl_2_, 5 mM KCl, 2 mM MES, pH 5.7), followed by washing with ½ MS or W5 solution for three times before imaged with the Leica SP8 confocal microscope. Images were captured at 552 nm laser excitation and 570-620 nm emission. The pinhole was set at one Airy unit, and the imaging processing was carried out by using the Leica Application Suite X (LAS X) software.

### MAPK assay

Ten-day-old seedlings grown vertically on ½ MS plates were transferred to ddH_2_O overnight for recovery. Then, ddH_2_O was replaced by 100 nM peptides for the indicated time. Each sample containing three seedlings was grounded in 40 μL of extraction buffer (150 mM NaCl, 50 mM Tris-HCl, pH 7.5, 5 mM EDTA, 1% [v/v] Triton X-100, 1 mM Na3VO4, 1 mM NaF, 1 mM DTT, 1:200 complete protease inhibitor cocktail from Sigma). The supernatant was collected after 13, 000 g centrifugation for 5 min at 4 °C and protein samples with 1 x SDS buffer were loaded on 10% (v/v) SDS-PAGE gel before transfer to a PVDF membrane, which was then blotted using α-pERK1/2 antibodies (Cell Signaling, Cat # 9101) for the detection of phosphorylated MPK3, MPK4, and MPK6.

### BIK1 phosphorylation assay

WT protoplasts were transfected with *pHBT-35S::BIK1-HA* for 12 hr followed by treatment with 1 μM SCOOP peptides for 15 min. Total proteins were isolated with extraction buffer (150 mM NaCl, 50 mM Tris-HCl, pH 7.5, 5 mM EDTA, 1% Triton X-100, 1 mM Na3VO4, 1 mM NaF, 1 mM DTT, 1:200 complete protease inhibitor cocktail from Sigma). The supernatant collected after 13, 000 g centrifugation for 5 min at 4 °C were loaded on 10% SDS-PAGE gel before transfer to a PVDF membrane, which was then blotted with α-HA antibodies (Roche, Cat # 12013819001).

### Co-immunoprecipitation (co-IP) assay

For co-IP assays in protoplasts, protoplasts were co-transfected with *pHBT-35S::MIK2-HA* and *pHBT-35S::BAK1-FALG* or *pHBT-35S::SERK4-FLAG* at 50 μg DNA for 500 μL protoplasts at the density of 2 × 10^5^/mL for each sample and were incubated for 12 hr. After treatment with 1 μM SCOOP peptides for 15 min, samples were collected by centrifugation at 200 g for two min and lysed in 300 μL IP buffer (20 mM Tris-HCl, pH7.5, 100 mM NaCl, 1 mM EDTA, 2 mM DTT, 10% [v/v] Glycerol, 0.5% [v/v] Triton X-100 and protease inhibitor cocktail from Sigma) by vortexing. After centrifugation at 10, 000 g for 10 min at 4 °C, 30 μL of supernatant was collected for input controls. The remaining supernatant was pre-incubated with protein-G-agarose beads at 4 °C for 1 hr with gentle shaking at 100 g on a rocker. IP was carried out with α-FLAG agarose for 3 hr at 4 °C. Beads were collected by 500 g centrifugation for 2 min and washed three times with washing buffer (20 mM Tris-HCl, pH7.5, 100 mM NaCl, 1 mM EDTA, 0.1% [v/v] Triton X-100) and once with 50 mM Tris-HCl, pH7.5. Immunoprecipitated proteins and input proteins were analyzed by immunoblotting with α-HA or α-FLAG antibodies (Sigma-Aldrich, Cat # A8592).

For co-IP assays in transgenic seedlings, leaves of four-week-old *pMIK2::MIK2-GFP* transgenic plants were hand-inoculated with 1 μM SCOOP10^B^ or SCOOP12 and were treated for 30 min, followed by total protein isolation with extraction buffer (50 mM Tris-HCl, pH 7.5, 150 mM NaCl, 1% [v/v] NP-40 and 1:200 protease inhibitor cocktail from Sigma). Co-IP was carried out the same as for co-IP with protoplasts except that GFP-Trap agarose beads (Chromotek) were used instead. Immunoprecipitated proteins and input proteins were analyzed by immunoblotting with α-GFP (Roche, Cat # 11814460001) or α-BAK1 antibodies ^55^.

### Protein expression and purification

The ECD domains of MIK2 (AA residues 1-707) and BAK1 (AA residues 1-220) fused with a six-HIS tag at the C-terminus were expressed using the Bac-to-Bac baculovirus expression system (Invitrogen) in High Five cells at 22 °C as reported previously ^23^. Secreted MIK2^ECD^-6xHIS and BAK1^ECD^-6xHIS proteins were purified using Ni-NTA (Novagen) and size-exclusion chromatography (Hiload 200, GE Healthcare) in buffer containing 10 mM Bis-Tris pH 6.0 and 100 mM NaCl.

### Isothermal titration calorimetry (ITC) assays

The binding affinities of MIK2^ECD^ with SCOOP12 or SCOOP12^SS/AA^ peptides were measured on a MicroCalorimeter ITC200 (Microcal LLC) at 25 °C as described previously ^23^. SCOOP12 or SCOOP12^SS/AA^ peptides (600 μM) were injected (1.5 μL per injection) into the stirred calorimeter cells containing 30 μM MIK2^ECD^. ITC data were analyzed using MicroCal Origin 7.0.

### Surface plasmon resonance (SPR) assays

The binding kinetics and affinities of MIK2^ECD^ with SCOOP12 or SCOOP12^SS/AA^ peptides were assessed on a Biacore T200 instrument (GE Healthcare, Pittsburgh, PA) with CM5 sensor chips. The MIK2^ECD^ proteins were exchanged to 10 mM sodium acetate (pH 5.5), and the peptides used for SPR was dissolved in HBS-EP+ (10 mM HEPES, 150 mM NaCl, 3 mM EDTA and 0.05% [v/v] surfactant P20 (GE Healthcare)). About 4700 response units of MIK2^ECD^ proteins were immobilized on a CM5 chip, and a blank channel was used as a negative control. SCOOP12 or SCOOP12^SS/AA^ peptides were diluted into indicated concentrations and injected at a flow rate of 30 μL min^−1^ for 2 min, followed by dissociation for 5 min. After dissociation, 5 mM NaOH was injected for 30 s to remove any non-covalently bound proteins from the chip surface. The binding kinetics was analyzed with the software Biaevaluation^®^ Version 4.1 using the 1:1 Langmuir binding model.

### Gel filtration assays

The purified MIK2^ECD^ proteins (0.5 mg for each) and BAK1^ECD^ proteins (0.2 mg for each) were incubated together with SCOOP12 or SCOOP12^SS/AA^ (0.1 mg for each) on ice for 1 hr. Then, each of the mixtures was separated by gel filtration (Hiload 200, GE Healthcare), and the peak fractions were separated on SDS-PAGE followed by Coomassie blue (CBB) staining.

### Phylogenetic analysis and graphical illustration

Protein sequences were retrieved from the NCBI database. Multiple sequence alignments were generated with ClustalW^56^. Phylogenetic analysis was carried out using MEGAX with the neighbor-joining method with 1000 bootstrap replicates^57^. The phylogenetic tree was visualized in an interactive tree of life (iTOL, https://itol.embl.de/). Homology modeling and visualization of *F. graminearum* protein FGSG_07177 were performed with Pymol (The PyMOL Molecular Graphics System, Version 1.2r3pre, Schrödinger, LLC.).

### Statistical analysis

Statistical analyses were all done in Excel with built-in formulas. The *P*-values were calculated with two-tailed Student’s unpaired *t*-test analysis for binary comparison or one-way ANOVA and Tukey’s post hoc honest significance test to compare more than two genotypes or treatments. The measurements shown in box plots display the first and third quartiles and split by medians (center lines), with whiskers extending to 1.5-fold the interquartile range from the 25th and 75th percentiles. Dots represent outliers. Asterisks illustrate the *P* values: ***, *P* < 0.001; **, *P* < 0.01 and *, *P* < 0.05.

## ACKNOWLEDGMENTS

We thank the *Arabidopsis* Biological Resource Center (ABRC) for providing the *Arabidopsis* T-DNA insertion lines, Dr. Reidunn B. Aalen (University of Oslo, Norway), Dr. Zhi Qi (Inner Mongolia University, China), Dr. Frederick M. Ausubel (Harvard Medical School, US), and Dr. Marc Knight (Durham University, UK) for providing Arabidopsis mutants or transgenic lines, and Dr. Weicai Yang (IGDB, CAS) for providing *MIK2-HA* construct. We also thank Dr. Ralph Hückelhoven (Technical University of Munich, Germany) for discussions, Dr. Jinrong Xu (Purdue University, US), and Dr. Huiquan Liu (Northwest A&F University, China) for assisting on the *Fusarium* genome sequence analysis, Dr. Wei Zhang (Shandong University, China) for technique supports, and members of the laboratories of L.S. and P.H. for discussions and comments of the experiments. The work was supported by National Science Foundation (NSF) (IOS-1951094) and NIH (R01GM092893) to P.H., NIH (R01GM097247), the Robert A. Welch Foundation (A-1795) to L.S., National Natural Science Foundation of China (NSFC) to S. Hou (31500971) and Z. H (31971119), and Youth Innovation Technology Project of Higher School in Shandong Province and to S. Hou (2020KJF013) and R. M (2019KJD003). The authors have declared no conflict-of-interest.

## AUTHOR CONTRIBUTIONS

S. H., D. Liu., P.H., and L.S conceived, designed experiments, analyzed data, and wrote the manuscript; S. Hou identified SCOOP and SCOOP-LIKE sequences, generated Arabidopsis transgenic and *mik2* mutant lines, performed physiological analysis, RNA-seq and RT-qPCR analyses, SPR assay, ROS production assay, and cytosolic calcium assay; D. Liu performed immunoblotting analysis, protein-protein interaction analysis, ethylene production measurement, microscopy assays, homology modeling, and statistical analysis; S. Huang under the supervision of Z.H. and J.C. purified MIK2^ECD^ and BAK1^ECD^ proteins and performed ITC and gel filtration analyses; D. Luo contributed to the root growth and ROS production assays, generated *pMIK2::MIK2-GFP/mik2-1* transgenic lines; Z. L. performed phylogenetic analysis, sequence alignment analysis, the *Fusarium* genome sequence analysis, and partial SPR assay; P. W analyzed pathogen SCOOPLs. R.M. analyzed the data.

---

**Figure 1.**
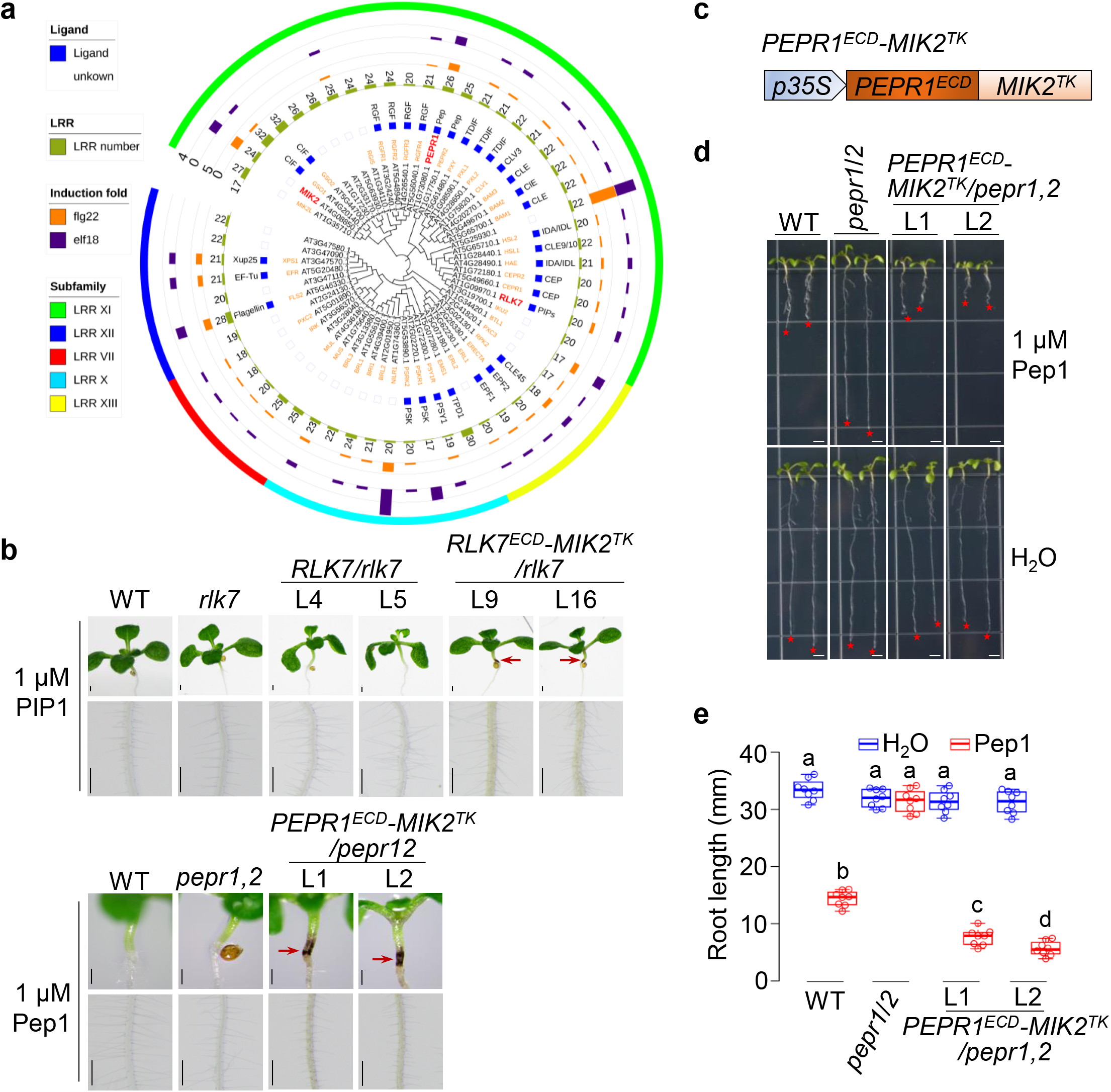
The cytosolic kinase domain of MIK2 triggers specific responses. **a.** *MIK2, RLK7,* and *PEPR1* are upregulated upon flg22 and elf18 treatments and phylogenetically related. Phylogenetic analysis and MAMP-induced expression of 56 LRR-RKs from the subfamily X (light blue curved line), XI (green curved line), XII (blue curved line), XIII (yellow curved line), and VII (red curved line). The LRR-RK full-length sequences were retrieved from NCBI (https://www.ncbi.nlm.nih.gov/) for MEGAX phylogenetic analysis using the neighbor-joining method with 1000 bootstrap replicates. The phylogenetic tree was displayed by iTOL v5 online software (https://itol.embl.de/). Blue squares indicate the cognate known ligands. Green bars with numbers indicate the number of LRRs for the corresponding LRR-RKs. Orange and purple bars indicate the induction levels of the cognate LRR-RK genes upon flg22 or elf18 treatment, respectively, according to the data from GENEVESTIGATOR V3. MIK2, RLK7, and PEPR1, used in this study, were highlighted in red. **b.** Activation of the cytosolic kinase domain of MIK2 induces the brown roots and darkened hypocotyl-root junctions. Transgenic seedlings of *RLK7^ECD^-MIK2^TK^*/*rlk7,* but not WT or *RLK7/rlk7*, show the brown roots and darkened hypocotyl-root junctions upon PIP1 treatment (top panel). *PEPR1^ECD^-MIK2^TK^/pepr1,2* transgenic seedlings show similar phenotypes upon Pep1 treatment (bottom panel). Seedlings of WT (Col-0), *rlk7*, two representative lines (L9 and L16) of *RLK7^ECD^-MIK2^TK^*/*rlk7*, two lines (L4 and L5) of *RLK7/rlk7*, *pepr1,2* mutant, and two representative lines (L1 and L2) of *PEPR1^ECD^-MIK2^TK^/pepr1,2* were grown on ½ MS plates with or without 1 μM PIP1 (upper) or 1 μM Pep1 (lower) for ten days. Red arrows indicate the hypocotyl-root junctions. Scale bar, 1 mm. **c.** Diagram of the *PEPR1^ECD^-MIK2^TK^* chimeric receptor. The extracellular domain of *PEPR1* (*PEPR1^ECD^*) was fused with the transmembrane and cytoplasmic kinase domain of *MIK2* (*MIK2^TK^*) to generate the *PEPR1^ECD^-MIK2^TK^* chimeric receptor gene. The *PEPR1^ECD^-MIK2^TK^* transgene under the control of a *35S* promoter was transformed into the *pepr1,2* mutant. **d, e.** Activation of the cytosolic kinase domain of MIK2 inhibits root growth. Two representative lines (L1 and L2) of *PEPR1^ECD^-MIK2^TK^*/*pepr1,2* transgenic seedlings showed more severe root growth inhibition to Pep1 treatment than WT plants. Seedlings were grown on ½ MS plates with or without 1 μM Pep1 for ten days (**d**). Red stars indicate the root tips. Scale bar, 4 mm. Quantification data of seedling root length are shown as the overlay of the dot plot and means ± SEM (**e**). Different letters indicate a significant difference with others (*P*<0.05, One-way ANOVA followed by Tukey’s test, *n*≥12). The experiments were repeated three times with similar results.

**Figure 2.**
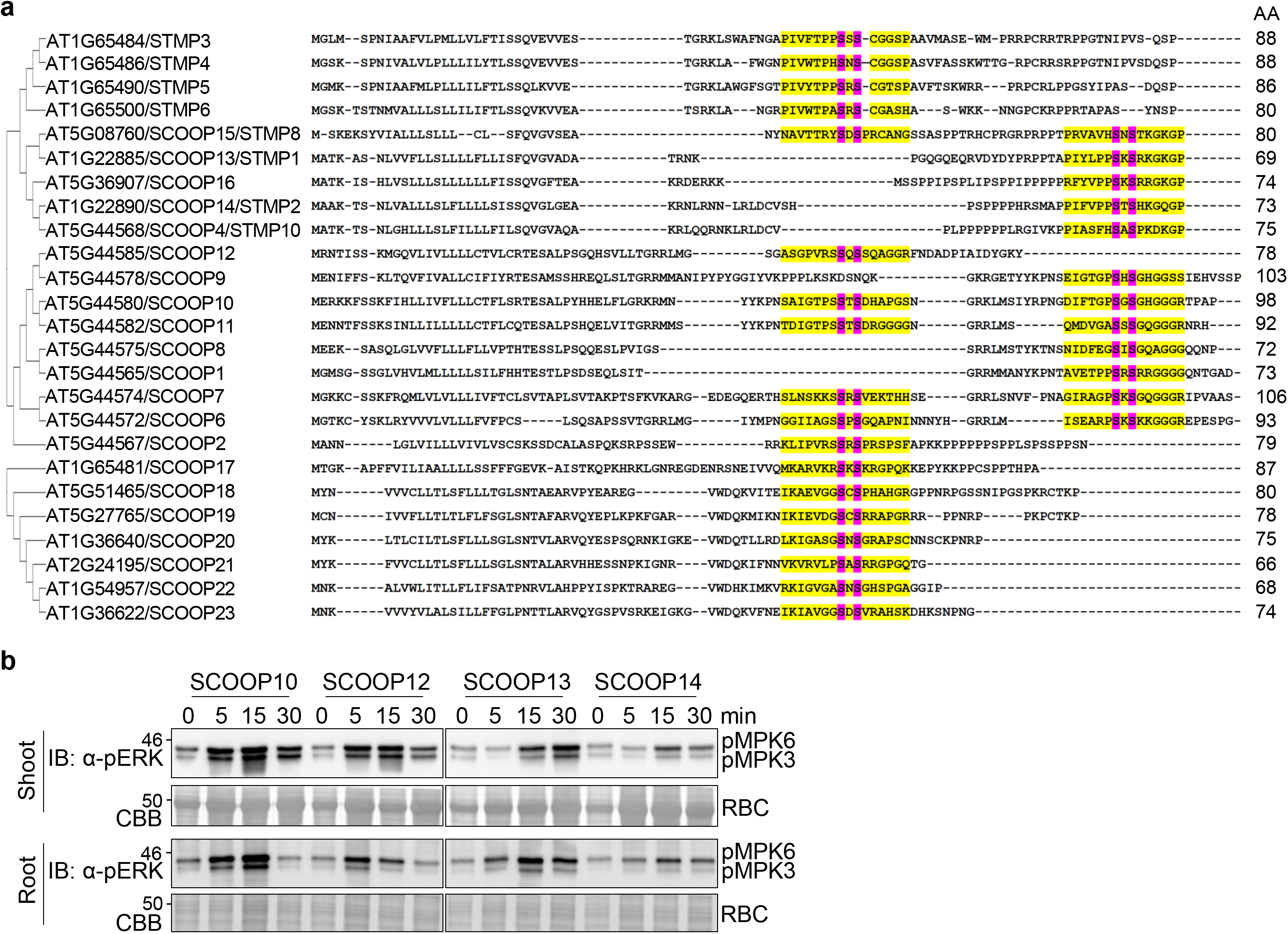
SCOOPs induce MAPK activation in shoots and roots. **a.** Sequence alignment and phylogenetic analysis of *Arabidopsis* SCOOP and STMP proteins. The amino acid sequences were aligned using ClustalW, and the phylogenetic tree was built by MEGAX using the neighbor-joining method with 1000 bootstrap replicates. The SxS motif and conserved serine residues are in yellow and pink, respectively. **b.** SCOOP peptides induce MAPK activation in shoots and roots. Ten-day-old WT seedlings grown in ½ MS liquid medium were treated with or without 1 μM peptides for the indicated time. Shoots and roots were collected separately for protein isolation and MAPK activation assays by immunoblotting with α-pERK antibodies (top panel), and the protein loading is shown by CBB staining for RBC (bottom panel).

**Figure 3.**
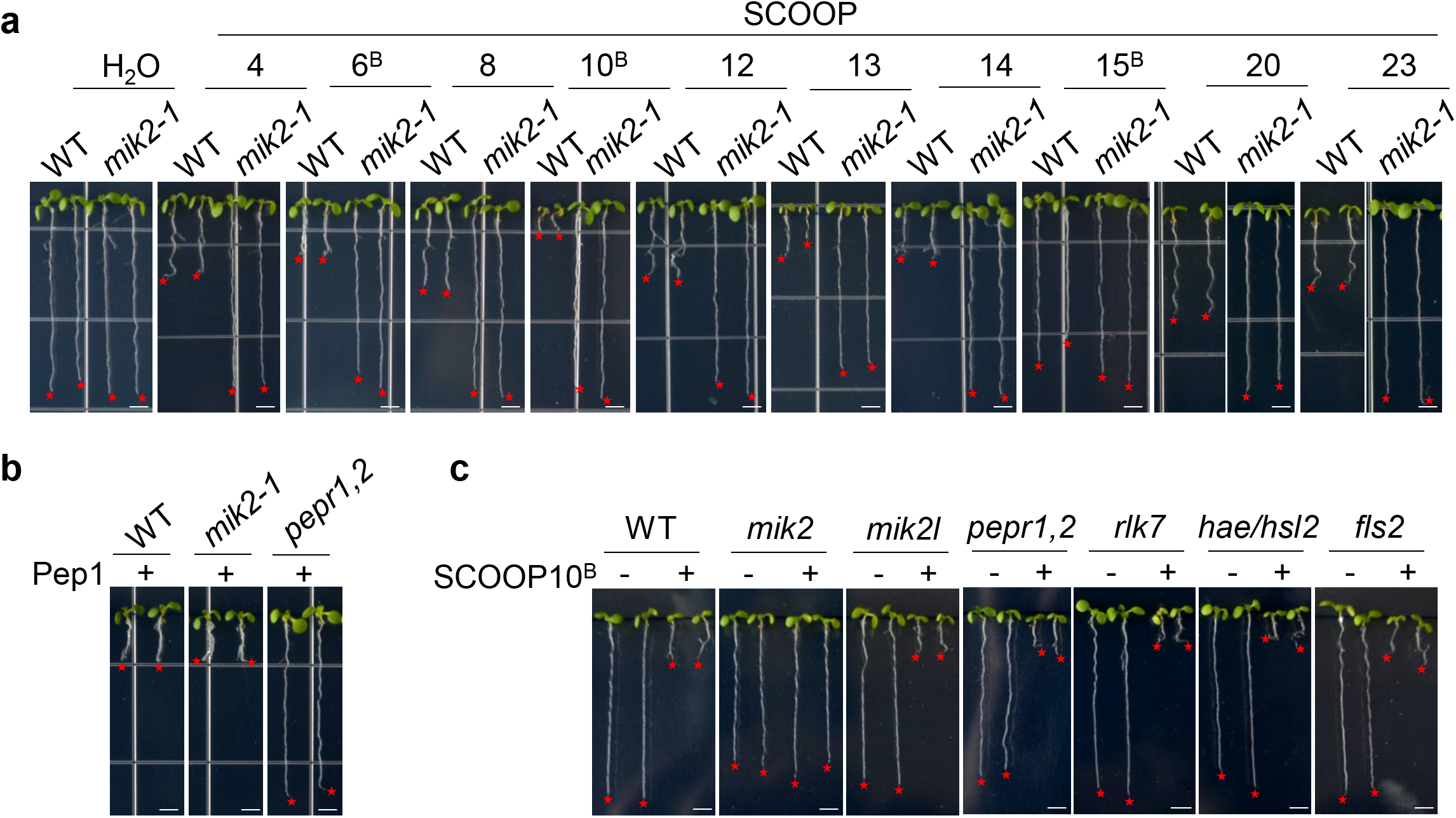
SCOOP-triggered root growth inhibition depends on MIK2, but not other related RKs. **a.** SCOOP-triggered root growth inhibition is blocked in the *mik2-1* mutant. WT and *mik2-1* mutant seedlings were grown on ½ MS plates with or without 1 μM SCOOP peptides for ten days. **b.** The *mik2-1* mutant is sensitive to Pep1 as WT. WT, *mik2-1*, and *pepr1,2* seedlings were grown on ½ MS plates with or without 1 μM Pep1 for ten days. **c.** SCOOP10^B^ peptides trigger robust root growth inhibition in *mik2l, fls2, pepr1/2, rlk7,* and *hae/hsl2* mutants. Seedlings were grown on ½ MS plates with or without 1 μM SCOOP10^B^ for ten days. Scale bar, 4 mm (**a-c**). The experiments were repeated three times with similar results.

**Figure 4.**
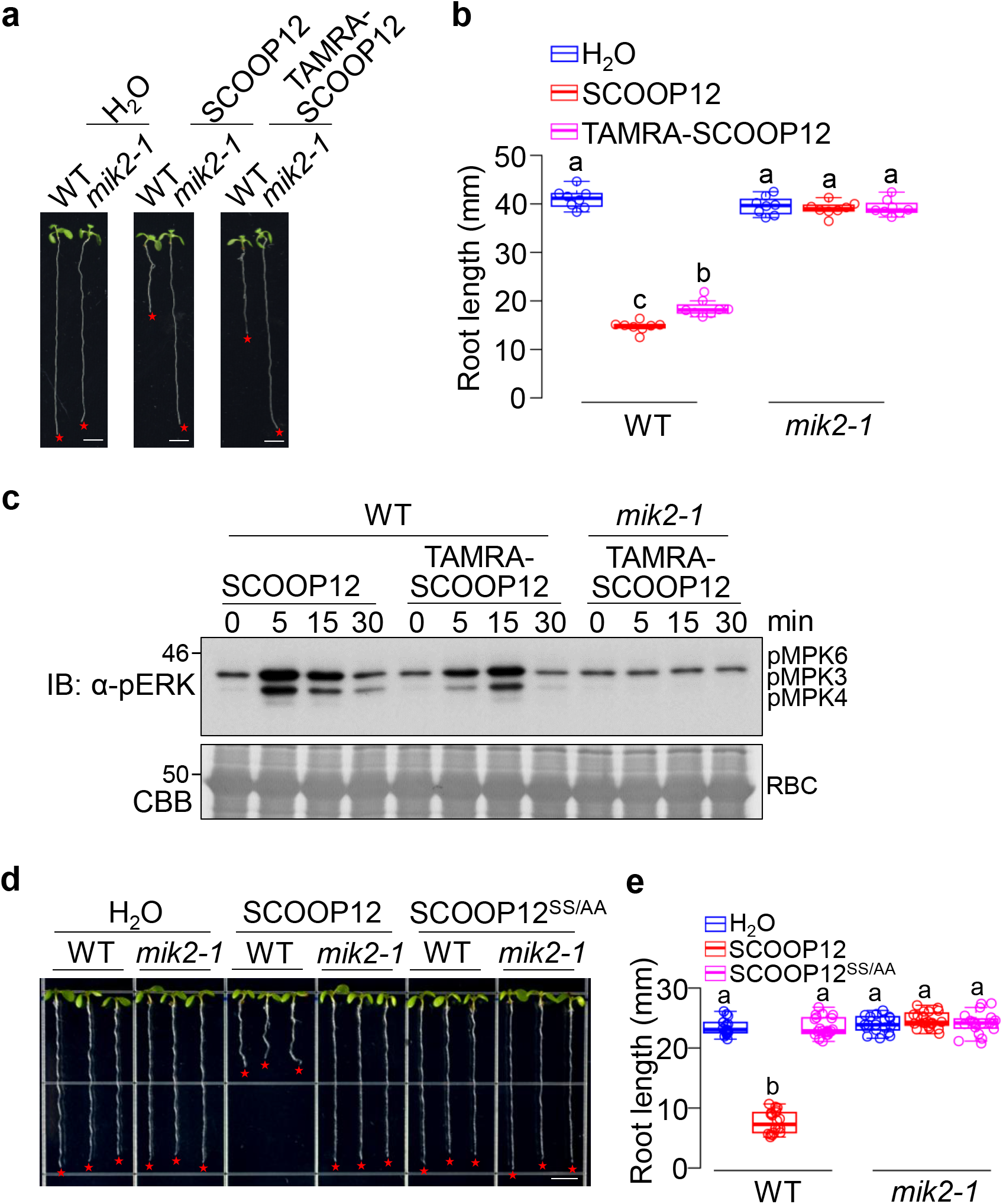
TAMRA-SCOOP12 triggers root growth inhibition in a MIK2-dependent manner. **a, b.** TAMRA-SCOOP12 peptides inhibit root growth in a MIK2-dependent manner. WT and *mik2-1* seedlings were grown on ½ MS plates with or without 1 μM TAMRA-SCOOP12 for ten days. Scale bar, 4 mm (**a**). Quantification data of seedling root length are shown as the overlay of dot plot and means ± SEM (**b**). Different letters indicate a significant difference with others (*P*<0.05, One-way ANOVA followed by Tukey’s test, *n*≥10). **c.** TAMRA-SCOOP12 peptides induce the MAPK activation in a MIK2-dependent manner. Ten-day-old WT and *mik2* seedlings grown in ½ MS liquid medium were treated with or without 1 μM SCOOP12 or TAMRA-SCOOP12 for the indicated time. The MAPK activation was analyzed by immunoblotting with α-pERK antibodies (top panel), and the protein loading is shown by CBB staining for RBC (bottom panel). **d, e** SCOOP12^SS/AA^ peptides are inactive to inhibit root growth. The assay and quantification were performed as in **a, b** with or without 1 μM SCOOP12 or SCOOP12^SS/AA^. Different letters indicate a significant difference with others (*P*<0.05, One-way ANOVA followed by Tukey’s test, *n*≥18). The experiments were repeated three times with similar results.

**Figure 5.**
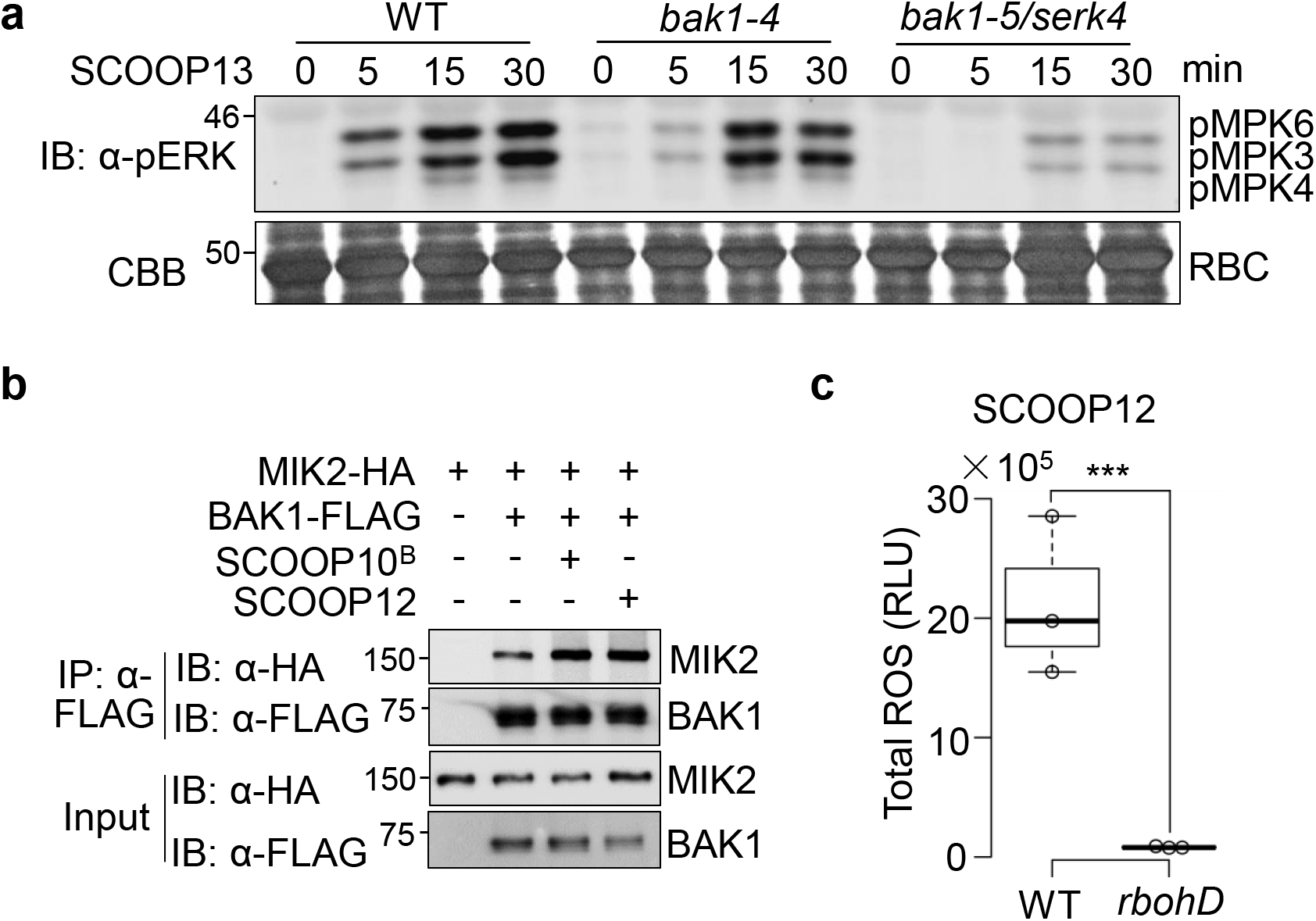
The SCOOP-triggered response depends on BAK1 and SERK4. **a.** SCOOP13-induced MAPK activation is compromised in *bak1-5/serk4.* Ten-day-old seedlings of WT, *bak1-4*, and *bak1-5/serk4* were treated with 100 nM SCOOP13 for the indicated time. The MAPK activation was analyzed by immunoblotting with a-pERK antibodies (top panel), and the protein loading is shown by CBB staining for RBC (bottom panel). **b.** SCOOPs enhance the association of MIK2 and BAK1 in protoplasts. Protoplasts from WT plants were co-transfected with HA-tagged MIK2 (MIK2-HA) and FLAG-tagged BAK1 (BAK1-FLAG), or a control vector (Ctrl) and incubated for 10 hr, followed by treatment with or without 1 μM SCOOP^B^ or SCOOP12 for 15 min. Proteins were immunoprecipitated with α-FLAG agarose beads (IP: α-FLAG), followed by immunoblotting (IB) with α-HA or α-FLAG antibodies (top two panels). IB on the bottom two panels shows the input controls before IP. **c.** SCOOP12-induced ROS production is blocked in the *rbohD* mutant. One-week-old seedlings grown on ½ MS plates were treated with 100 nM SCOOP12 for ROS measurement. Total luminescence counts as RLUs are shown as the overlay of the dot plot and means ± SEM (*** *P*<0.001, Student’s *t*-test, *n*=4). The above experiments were repeated three times with similar results.

**Figure 6.**
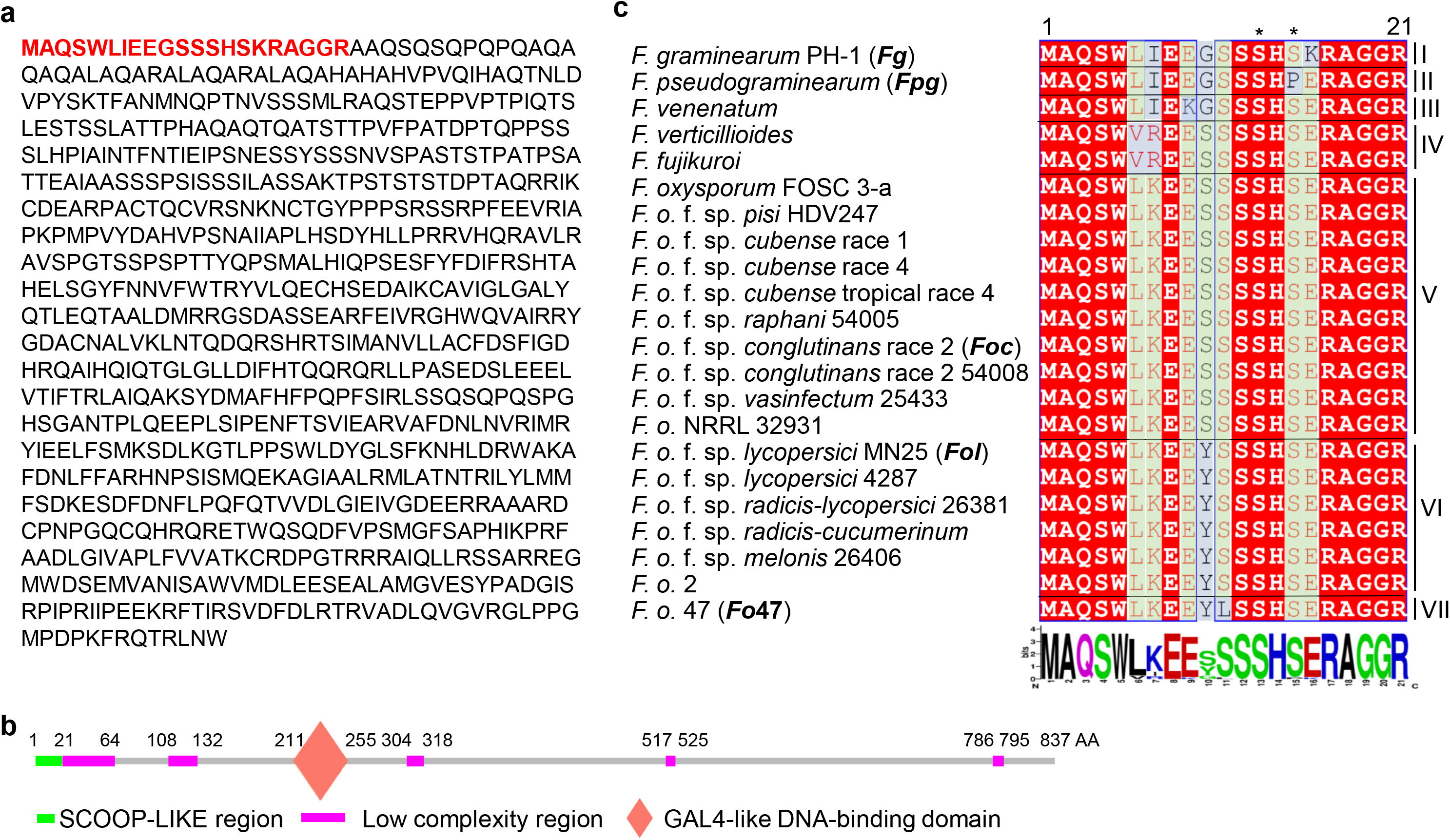
The FGSG_07177 protein from *F. graminearum* contains a SCOOP-LIKE (SCOOPL) sequence in its N-terminus. **a.** The full-length amino acid sequence of FGSG_07177. The sequence from *F. graminearum* was downloaded from the NCBI website, and the SCOOPL sequence was marked in red. **b.** Schematic diagrams of FGSG_07177 protein motifs. The predicated domains analyzed by the SMART database (https://smart.embl.de/) were highlighted with amino acid positions labeled on the top. **c.** SCOOPLs are conserved in *Fusarium* species. SCOOPL sequences were blast-searched in the NCBI database using *Arabidopsis* SCOOP10^B^ as the reference, aligned using ClustalW and visualized in ESPript3 (espript.ibcp.fr). The SxS motif was marked with stars on the top. The consensus of SCOOPs was conducted with WebLogo (http://weblogo.berkeley.edu/logo.cgi). SCOOPL peptides from *Fg*, *Fpg*, *Foc*, *Fol*, and *Fo*47 (highlighted in bold) were synthesized for bioassays.

**Figure 7.**
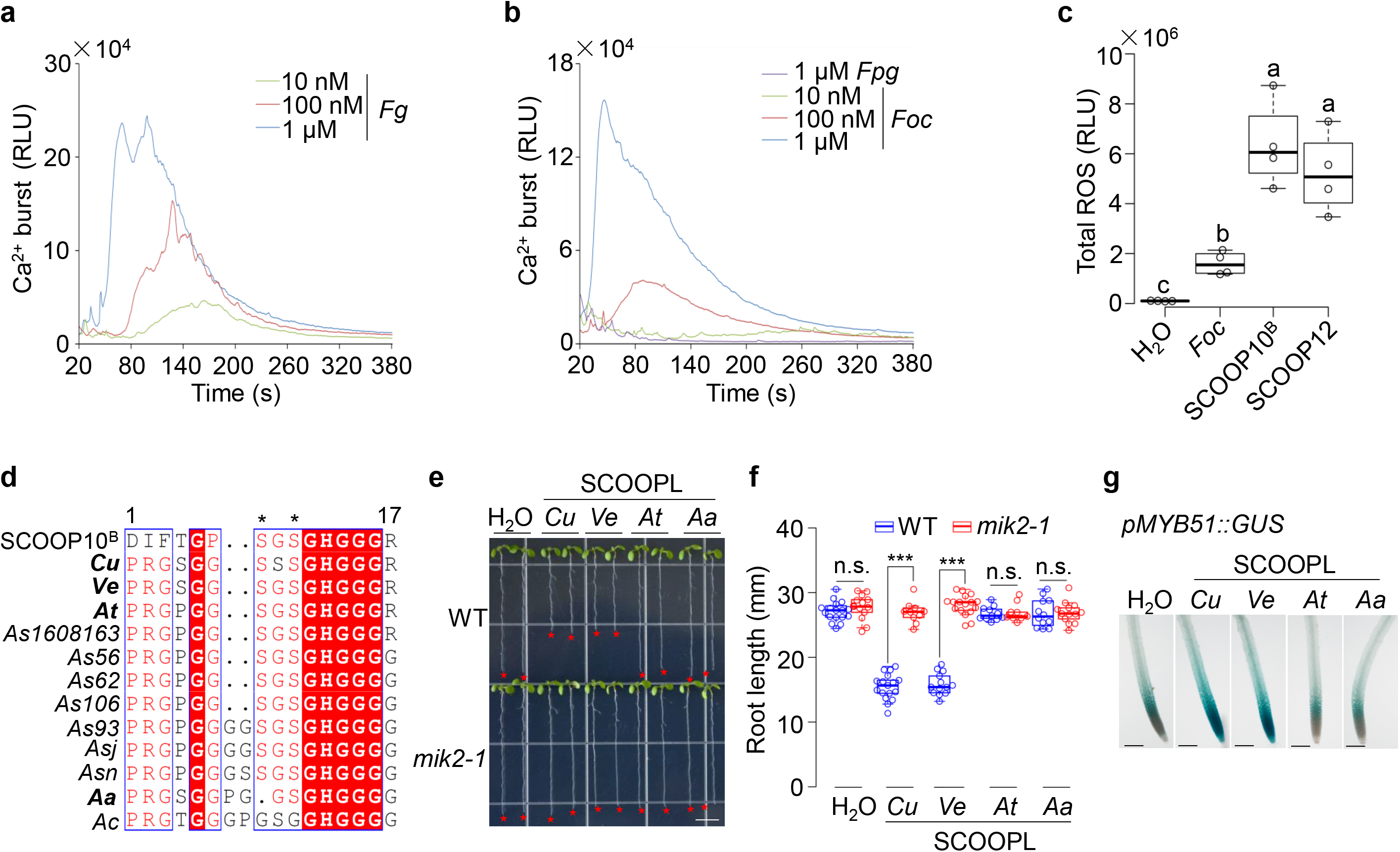
*Fusarium* and bacterial SCOOPLs induce immune responses. **a, b.** SCOOPLs from *Fg* and *Foc*, but not from *Fpg*, trigger the cytosolic Ca^2+^ increase. One-week-old transgenic seedlings expressing *p35S::Aquorin* grown on ½ MS plates were treated with or without peptides with the indicated concentrations for the continuous measurement of cytosolic Ca^2+^ concentration. **c.** *Foc*SCOOPL induces a weaker ROS production than SCOOP10^B^ and SCOOP12. One-week old WT seedlings were treated with 1 μM *Foc*SCOOPL, 100 nM SCOOP10^B^ or 100 nM SCOOP12 for ROS measurement. Total luminescence counts as RLUs are shown as the overlay of the dot plot and means ± SEM. Different letters indicate a significant difference with others (*P*<0.05, One-way ANOVA followed by Tukey’s test, *n*=4). **d.** Alignment of bacterial SCOOPL sequences. SCOOPL sequences from different bacterial species were retrieved by blast-searching the NCBI database using *Arabidopsis* SCOOP10^B^ as the reference, aligned using ClustalW, and visualized in ESPript3 (espript.ibcp.fr). The SxS motif was marked with stars on the top. SCOOPL peptides from *Cu, Ve, At,* and *Aa* (highlighted in bold fonts) were synthesized for bioassays. *Cu*, *Curvibacter* sp.; *Ve*, *Verminephrobacter eiseniae*; *At*, *Acidovorax temperans*; *A*1608163, *Acidovorax* sp. 1608163; *A*56, *Acidovorax* sp. 56; *A*62, *Acidovorax* sp. 62; *A*106, *Acidovorax* sp. 106; *A*93, *Acidovorax* sp. 93; *A*j, *Acidovorax* sp. JMULE5; *A*N, *Acidovorax* sp. NO-1; *Aa*, *Acidovorax avenae* subsp. *Avenae*; *Ac*, *Acidovorax citrulli*. **e, f.** Bacterial SCOOPLs inhibit root growth in a MIK2-dependent manner. WT and *mik2-1* mutant seedlings were grown on ½ MS plates with or without 1 μM peptides for ten days. Red stars indicate the root tips. Scale bar, 4 mm (**e).** Quantification data of seedling root length are shown as the overlay of the dot plot and means ± SEM (**f**) (*** *P*<0.001, n.s., no significant differences, Student’s *t*-test, *n*≥15). **g.** Bacterial SCOOPLs induce the expression of *pMYB51::GUS* expression in roots. One-week-old transgenic seedlings carrying *pMYB51::GUS* grown on ½ MS plates were treated with or without 1 μM bacterial SCOOPL peptides for 3 hr and subjected to GUS staining and distaining followed by photographing under a stereomicroscope. Scale bar, 1mm. The experiment was repeated three times with similar results.

**Supplementary Table 1. Primers used in this study.**

**Supplementary Table 2. Peptides used in this study.**

**Supplementary Table 3. Differentially expressed genes in WT Col-0 and *RLK7^ECD^-MIK2^TK^* transgenic plants upon PIP1 treatment.**

